# The ADAR1 editome reveals drivers of editing-specificity for ADAR1-isoforms

**DOI:** 10.1101/2021.11.24.469911

**Authors:** Renata Kleinova, Alina F. Leuchtenberger, Claudio Lo Giudice, Andrea Tanzer, Sophia Derdak, Ernesto Picardi, Michael F. Jantsch

## Abstract

Adenosine deaminase acting on RNA (ADAR) (also known as ADAR1) promotes A-to-I conversion in double-stranded and highly structured RNAs. ADAR1 has two isoforms transcribed from different promoters: ADAR1p150, which is mainly cytoplasmic and interferon-inducible, and constitutively expressed ADAR1p110 that is primarily localized in the nucleus.

Mutations in ADAR1 cause Aicardi – Goutières syndrome (AGS), a severe autoinflammatory disease in humans associated with aberrant IFN production. In mice, deletion of ADAR1 or selective knockout of the p150 isoform alone leads to embryonic lethality driven by overexpression of interferon-stimulated genes. This phenotype can be rescued by concurrent deletion of cytoplasmic dsRNA-sensor MDA5. These findings indicate that the interferon-inducible p150 isoform is indispensable and cannot be rescued by the ADAR1p110 isoform. Nevertheless, editing sites uniquely targeted by ADAR1p150 but also mechanisms of isoform-specificity remain elusive.

Here we combine RIP-seq on human cells expressing ADAR1 isoforms and combine this with analysis of isoform-specific editing patterns in genetically modified mouse cells to extensively investigate ADAR1-isoform binding- and editing characteristics.

Moreover, using mutated ADAR variants, we examine the effect of two unique features of ADAR1p150 on its target specificity: 1) cytoplasmic localization and 2) Z-DNA binding domain *α*. Our findings indicate that ZBD*α* contributes only minimally to p150 editing-specificity and that isoform-specific editing is directed mainly by the cytoplasmic localization of the editase.

## Introduction

Adenosine to inosine RNA editing (A-to-I RNA editing) is one of the major nucleotide modifications found in RNAs (Nishikura, 2016). In human and mouse transcriptomes, more than 10^6 and 10^5 A-to-I editing sites have been identified, respectively (Bazak et al., 2014a; Licht et al., 2019; Picardi et al., 2017). A-to-I editing occurs in structured or double-stranded regions of RNA. The majority of editing sites is located in interspersed nuclear elements (SINEs) that are frequently located in introns and 3’UTRs (Bazak *et al*., 2014a; Peng et al., 2012). When two or more repeats are found in opposite orientation within an RNA, they tend to basepair and therefore can produce the double-stranded structures required for RNA-editing (Bazak et al., 2014b). Cellular machineries typically interpret inosines as guanosines. Therefore, RNA editing affects many cellular processes, including protein-coding, RNA-splicing, -folding and turnover (Licht et al., 2016; Nishikura, 2016).

In mammals, RNA editing is mediated by two catalytically active adenosine deaminases acting on RNA (ADARs): ADAR1 (*Adar*) and ADAR2 (*Adarb1*). ADARs contain a conserved deaminase domain and two or three dsRNA-binding domains that bind to dsRNA and highly structured RNA (Figure 1 - figure supplement 1) (Nishikura, 2010). ADAR1 is expressed from two different promoters giving rise to two isoforms: I) interferon-inducible ADAR1p150 and II) constitutively expressed ADAR1p110 (Patterson and Samuel, 1995).

Dysfunctions of ADAR1 can lead to Aicardi-Goutières syndrome (AGS), a fatal childhood encephalopathy accompanied with aberrant IFN signature (Livingston et al., 2014; Rice et al., 2012). In mice, lack of ADAR1 leads to embryonic lethality by the embryonic day E12.5 that is manifested by IFN overproduction, hematopoietic failure, liver disintegration and widespread apoptosis (Hartner et al., 2004; Wang et al., 2004). Mice bearing a catalytically inactive ADAR1^E861A^ knock-in mutation die at E13.5, which indicates that A-to-I editing is indeed the essential function of ADAR1 (Liddicoat et al., 2015).

Although the lack of ADAR2 is also not tolerated in mice and leads to postnatal lethality due to progressive seizures, the phenotype can be elegantly rescued by inserting a single point mutation mimicking editing in the glutamate receptor Gria2 (Higuchi et al., 2000).

No such analogous ADAR1-substrate that could rescue the deleterious effect of ADAR1 dysfunction has been identified. Nevertheless, embryonic death of ADAR1-deficient mice can be rescued by concurrent deletion of the cytoplasmic dsRNA-sensor MDA5 or its downstream adaptor protein MAVS (Figure 1 - figure supplement 2) (Liddicoat *et al*., 2015; Mannion et al., 2014; Pestal et al., 2015). Importantly, the unique deletion of the ADAR1p150 isoform is embryonic lethal with IFN-overproduction and can be rescued in the same manner as the full ADAR1 knockout – by additional deletion of MDA5 (Pestal *et al*., 2015). Moreover, a specific knockout of ADAR1p110 in mice is viable without aberrant IFN signature (Kim et al., 2021).

Taken together, this shows that ADAR1p150–mediated editing is a specific and essential regulator of the MDA5-MAVS pathway and that ADAR1p110 alone cannot prevent an undesired innate immune response (Kim *et al*., 2021; Pestal *et al*., 2015). ADAR1p150 contains a unique N-terminus bearing Z-DNA binding domain *α* (ZBD*α*) and a nuclear export signal (NES) that mediates a prevalently cytoplasmic localization. In contrast, ADAR1p110 is localized in the nucleus. However, both isoforms can shuttle between the nucleus and the cytoplasm, which is more prominent under certain circumstances such as viral infection (Barraud et al., 2014; Eckmann et al., 2001; Poulsen et al., 2001; Strehblow et al., 2002). ZBD*α* binds the unusual left-handed conformation of DNA and RNA (Brown et al., 2000). However, whether the ZBDs contribute to substrate binding or editing-specificity of ADAR1p150 is not fully understood. One of the most common ADAR1 mutations in AGS patients, P193A, lies in ZBD*α*. In AGS patients, the ADAR1^P193A^ allele exists in a compound-heterozygous stage with a second dysfunctional ADAR1 allele (Crow et al., 2015; Rice *et al*., 2012).

Interestingly, mice with mutations introduced in the ZBD*α* that extensively affect binding to nucleic acids in Z-conformation (N175A+Y179A, W197A) display reduced editing of SINE containing RNAs. Moreover, these mice also developed a spontaneous MAVS-depended IFN- response (de Reuver et al., 2021; Nakahama et al., 2021; Tang et al., 2021). Still, mice carrying homozygous mutations P195A (mimicking human P193A) in ZBD! were phenotypically normal. Most interestingly, the human AGS genotype was recapitulated once compound heterozygous mice were generated by introducing a copy of the ADAR1^P195A^ mutant allele to a second ADAR1^-^ or ADAR1p150^-^ deleted allele. The mutant mice developed a severe disease that was driven by MDA5 (Maurano et al., 2021b). Considering the lack of phenotype in homozygous ADAR1^P195A^ mutant mice and the absence of AGS patients carrying homozygous ZBD*α* mutations it seems that this domain is not a main driver of AGS but rather contributes to the disease. Moreover, the contribution of ZBD*α* to ADAR1p150 editing-specificity and on the functions of nuclear and cytoplasmic ADAR1p150-editing remain elusive.

In our study, we employ a combination of RIP-seq on human cells and isoform-specific editing pattern analysis in mouse cells to extensively examine ADAR1-isoform binding- and editing characteristics. We collect and comprehensively analyze a high-quality ADAR1p150- and ADAR1p110-specific mouse-editome. Moreover, using mutated ADAR versions, we investigated the role of cytoplasmic localization and of ZBD*α* on ADAR1p150 editing-specificity. Our findings suggest that ZBD*α* has a minor contribution to editing specificity and that ADAR1p150 editing patterns are mostly driven by the cytoplasmic localization of this protein isoform.

## Results

To determine the target specificity of ADAR1 isoforms, we employed two different approaches: on the one hand, we identified interacting RNAs in HEK 293 cells transfected with ADAR1 isoforms using RIP-seq. On the other hand, we identified RNAs that can become edited by ADAR1 isoforms in MEF cells. To do so, ADAR1 deficient MEFs were transfected with ADAR1 isoforms, and editing patterns were determined. Jointly, the two approaches give a robust overview of the ADAR1-editome.

Further, we determined the impact of distinct motifs altering the localization and nucleic acid binding properties of ADAR1-isoforms to assess their contribution to isoform-specific RNA-targeting.

### Methylene blue-enhanced ADAR1-RIP effectively captures specific targets

To examine the *in vivo* interaction between ADAR1 and its substrates, we employed RNA immunoprecipitation and sequencing (RIP-seq). To determine optimal RIP-seq conditions, we compared the original RIP protocol (Zambelli and Pavesi, 2015) with crosslink-RIP using either formaldehyde (fRIP) or methylene blue (MB) photo-crosslinking (Figure 1 - figure supplement 3) (Liu et al., 1996; Ricci et al., 2014). MB intercalates in dsRNAs, opens the helix, and thus facilitates cross-linking between double-stranded RNA-binding proteins (RBPs) and RNA in the presence of visible light. Moreover, we applied additional cross-linking with UV at 254 nm to achieve even stronger ADAR1-RNA binding (Lin and Miles, 2019). All examined protocols were first tested only with the ADAR1p150-isoform.

To compare the efficiency of the used protocols, we determined the fold-enrichment of known ADAR1-substrates in IP-fractions relative to corresponding inputs. Mock_IPs were used as controls. Primers were designed to span exonic editing site (ES) in Azin1 (chr8:103841636), a non-repetitive ES in the 3’UTR of Pum2 (chr2: 20450819), and the heavily edited 3’UTR of Nicn1. We observed a 10-20 fold enrichment of all 3 test substrates in MB+UV RIP while the other tested conditions only showed moderate, 2-3 fold enrichment (Figure 1 - figure supplement 4). Thus, our data show that MB+UV is indeed a powerful cross-linking method to detect ADAR1-RNA interactions.

### MB+UV RIP-seq identifies a distinct binding distribution of ADAR1-isoforms

Next, MB-UV RIP-seq was performed for ADAR1p150 and ADAR1p110 isoforms in triplicates. As a control, MOCK transfected cells were processed in parallel (in duplicates).

As expected, control IPs with untransfected samples showed no enrichment of ADAR1- specific substrates (Figure 1 - figure supplement 5). ADAR1p110-IP, when compared to ADAR1p150-IP, displayed only mild enrichment of selected targets. This indicates that the examined targets are prevalently bound and likely edited by ADAR1p150, a notion that was confirmed for Azin1 (chr8:103841636) by Sanger sequencing (Figure 1 - figure supplement 5). We prepared triplicate libraries from ribo-minus selected RIP-samples. We obtained between 6 to 13 million uniquely mapped 75 bp paired-end reads (Figure 1 – table 1). Next, peaks were called for these mapped reads to identify ADAR1-isoform binding substrates and regions.

Peaks that also appeared in the empty-transfection control were subtracted from ADAR1p150 and ADAR1p110 peaks to exclude unspecific peaks. This resulted in 14,191 peaks for ADAR1p150 and 2,921 peaks for ADAR1p110. Additionally, we excluded peaks shorter than 40 nt. Peak size is based on genomic rather than transcriptomic coordinates, thus ADAR1p150 peaks appear longer (the median size of ADAR1p150 peak is 2105 nt while ADAR1p110 peaks are 524 nt on average) (Figure 1 - figure supplement 6). This is most likely caused by cytoplasmically localized ADAR1p150 mostly binding to exonic regions. Consequently, these peaks map across introns. We, therefore, calculated the median width of ADAR1p150 and ADAR1p110 peaks and excluded all peaks longer than 3× the median peak size to omit extremely long peaks. Using all the above-mentioned filtering criteria, we identified 10,255 peaks for ADAR1p150 and 2,289 peaks for ADAR1p110 (Figure 1A) (Figure1 – table 2). The considerable discrepancy in the identified peak numbers can likely be explained by the higher reaction dynamics of ADAR1p110 editing which occurs mostly co-transcriptionally in the nucleus (Bentley, 2014; Licht *et al*., 2016). Thus, detecting RNAs transiently bound by ADAR1p110 might be less efficient than RNAs bound by cytoplasmic ADAR1p150 which likely will be bound longer. The majority of ADAR1p110 peaks is located in introns which corresponds well with its predominantly nuclear localization. In contrast, ADAR1p150 peaks largely fall into exonic regions, including a large fraction of 3’ UTRs (Figure 1B). A quite prominent portion of intronic ADAR1p150 peaks is probably caused by the annotation of peaks to genomic coordinates. Therefore, spanned introns also appear in the final peak annotation, although they are not physically present in the peaks (Figure 1-figure supplement 7). The ADAR1p150- and ADAR1p110 peak-set showed a limited overlap of 342 peaks for ADAR1p110, 308 peaks for ADAR1p150, respectively (ADAR1p150 peaks overlap with more than 1 ADAR1p110 peak) (Figure 1C). Thus about 12% of all ADAR1p110 peaks overlap with ∼2% of all ADAR1p150 peaks. This modest overlap is presumably caused by the limited binding of ADAR1p150 to intronic regions compared to ADAR1-p110, which leads to a shifted distribution of peaks for these two isoforms (Figure 1D). Almost 50% of ADAR1p150-peaks span non-repetitive regions, whereas only 14% of ADAR1p110-peaks lie in non-repetitive regions (Figure 1E). This observation greatly correlates with the gene-region distribution of the peaks. Whilst ADAR1p110 binds mostly introns that contain a high portion of repeats, ADAR1p150 also binds in 3’UTRs and exons. The majority of the interacting repeats accounts for SINE elements in the case of both isoforms. However, ADAR1p150-peaks span also a significant portion of simple repeats, LINE and DNA transposons (Figure 1F). In summary, our RIP seq experiments identified distinct binding regions for ADAR1-isoforms. While ADAR1p110 binds mostly intronic regions, ADAR1p150 also binds to exons and 3’UTRs.

**Figure 1:**
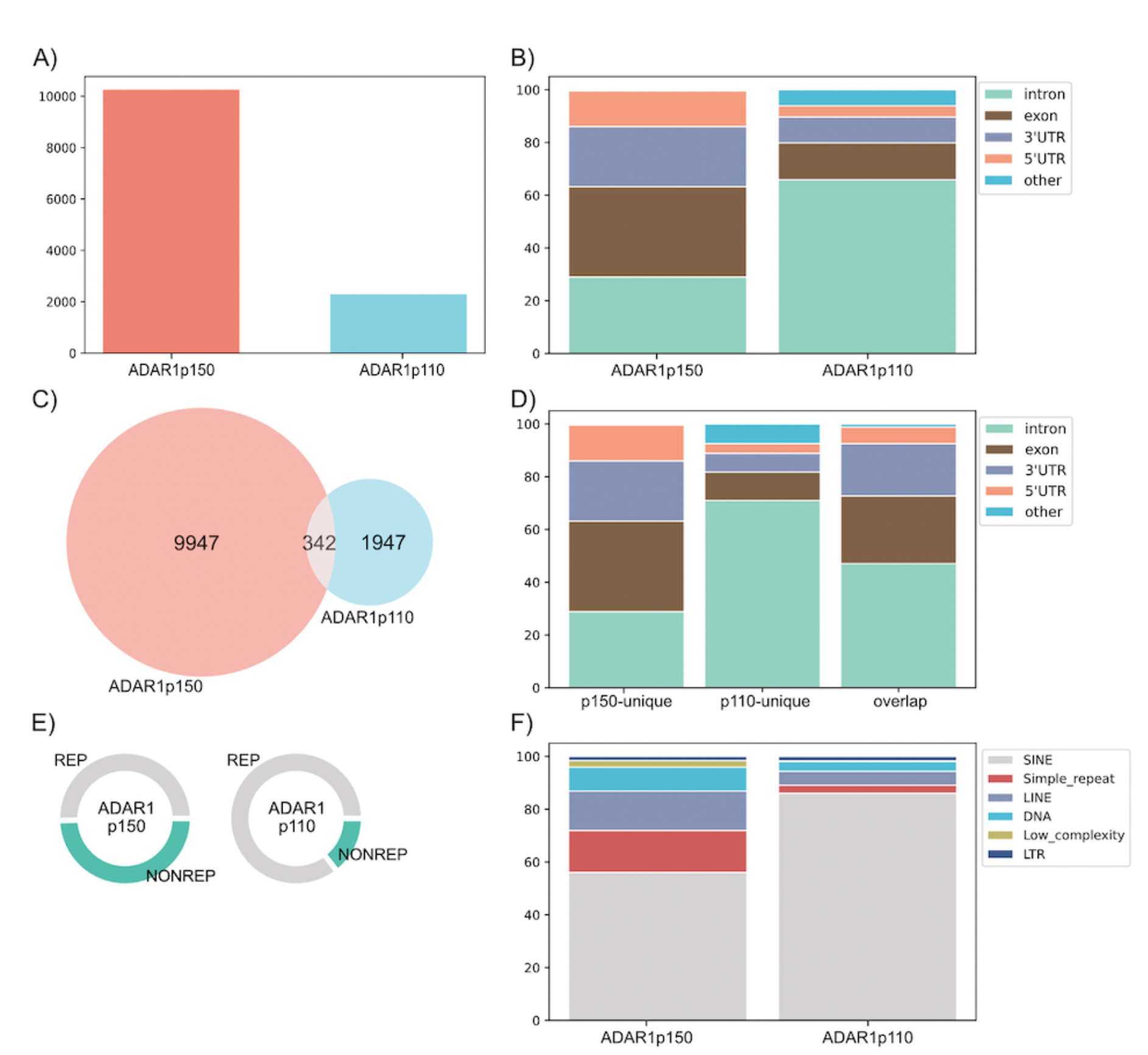
RIP-seq identifies distinct binding regions for ADAR1-isoforms. A) Number of peaks identified by peak-caller after passing filtering criteria. ADAR1p150: 10,255 peaks, ADAR1p110: 2,289; B) Peak-distribution in gene regions. ADAR1p110 mainly binds in introns whereas ADAR1p150 shows more complex binding involving exons and 3’UTRs; C) Isoform-specific and overlapping peaks. (Depicted ADAR1p110 overlap - 342); D) Distribution of isoform-specific and overlapping peaks to gene regions. E) Peak distribution to repeats and non-repetitive regions. While only 14% of ADAR1p110- peaks cover unique regions almost half of ADAR1p150-peaks span non-repetitive regions. F) The majority of interacting repeats locates to SINE elements, especially in the case of ADAR1p110. ADAR1p150 binds more diverse repeats, including also simple repeats, LINE and DNA transposons.

### Restoring ADAR1 expression in A-to-I editing deficient cells

After having shown a distinct binding preference for ADAR1p150 and ADAR1p110 isoforms in human cells, we wanted to evaluate ADAR1 isoform-specific editing preferences. To do this in an experimentally clean system, we turned to mouse cells that are deficient in ADAR1 and ADAR2 and therefore showed no background editing (MEFs^ADAR1-/-; ADAR2-/-^). In these cells, we restored ADAR1 expression by electroporation of ADAR1p150, ADAR1p110, or RFP (editase negative control –1 replica) expressing plasmids (Figure 2 - figure supplement 1). Unless stated otherwise, all experiments were conducted in triplicates. Overexpressed ADAR1 isoforms showed their typical intracellular localization and distinct editing patterns (Figure 2 - figure supplement 2). 24 hours post electroporation, the cells were harvested, and total RNA was extracted. After ribo-minus depletion and library construction ≈ 40 mio paired-end reads were obtained per replica. RNA-editing at known sites was evaluated in each sample using REDItools (Lo Giudice et al., 2020; Picardi and Pesole, 2013).

**Figure 2:**
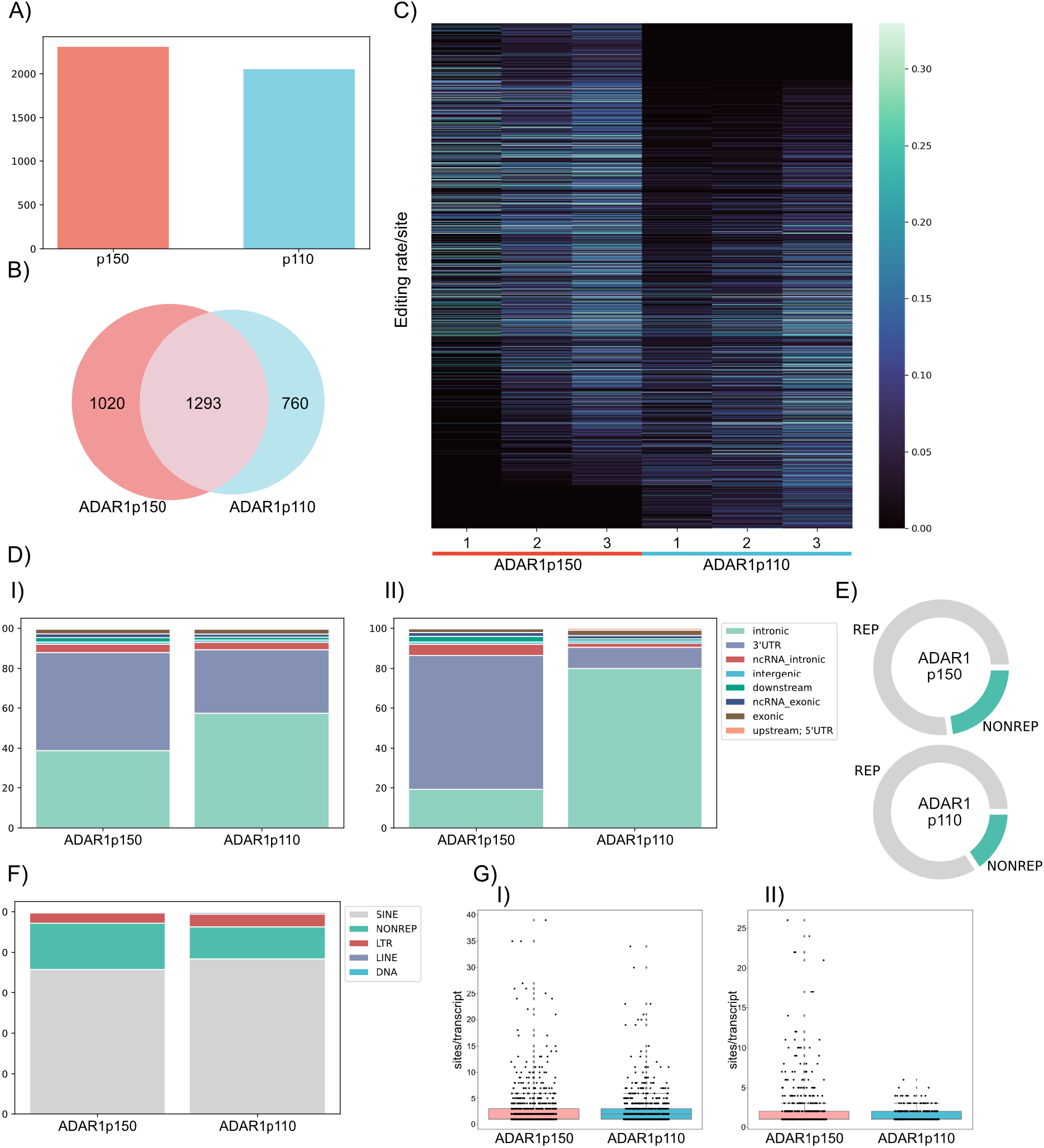
ADAR1p150 and ADAR1p110 display isoform-specific editing patterns. A) ≈ 3000 editing sites are efficiently edited (≥ 1 % editing ratio) by ADAR1p150 or ADAR1p110. B) ≈ 40% of detected sites are efficiently edited by both isoforms (1293); one third is preferentially edited by ADAR1p150 (1020), and 25% are specifically edited by ADAR1p110 (760). C) Heat map showing editing ratio of detected editing sites. Despite the large editing overlap between ADAR1-isoforms, the majority of sites still shows a clear preference for one or another isoform. D) Editing gene-region distribution of ADAR1- isoforms. I) Distribution of all efficiently edited sites. ADAR1p110 edits prevalently in introns. Apart from the intronic regions, ADAR1p150 efficiently edits a prominent portion of 3’UTRs. II) Distribution of editing sites mainly edited by ADAR1p150 or ADARp110: sites that are edited 2,5 × better by one isoform than by another. Editing sites edited by ADAR1p110 are almost exclusively intronic, whereas a major portion of ADAR1p150 specific sites locates to 3’UTRs. E) ADAR1p150 edits more nonrepetitive parts (∼23%) than ADAR1p110 (∼16%). F) The majority of editing locates to SINE elements. G) ADAR1p150 edits more hyperedited regions. I) Boxplot of all detected sites; II) Boxplot of sites mainly edited by ADAR1p150 or ADARp110: sites that are edited 2,5 × better by one isoform than by another; (y-axis: editing sites/transcript).

Whilst only negligible A to G transitions were found in RFP transfected cells (78 edited sites) (editase-negative control), editing raised remarkably upon transfection with any ADAR1 isoform (Figure 2 - figure supplement 3A). Noticeably less editing was detected in the 1st replica of ADAR1p150 because of the considerable rRNA contamination that led to overall low coverage and shallow editing detection (Figure 2 - figure supplement 3B). Still, when combining all samples and by considering all sites that were also covered by more than 5 reads in the negative control, we could identify 9.400 of more than 100.000 editing sites known in mice and stored in REDIportal (Licht *et al*., 2019; Picardi *et al*., 2017). Those sites were subjected to further filtering to acquire reliable sets of editing sites for each ADAR1- isoform. First, we excluded all sites also detected in the RFP transfection control. Next, we removed sites with low coverage (less than 5 reads per replicate) and editing rates <1%. Finally, only sites edited in 2 out of 3 replicas and sufficiently covered in both isoform-datasets (p150 and p110, average ≥ 10 reads) were subjected to further analysis (Figure 2 - figure supplement 3C). Following these stringent filtering criteria, we collected an authentic set of 3073 edited sites (Figure 2 - figure supplement 3D, Figure 2 – table 1).

### ADAR1p150 and p110 display specific editing patterns

A similar number of editing sites was detected for both ADAR1 isoforms. For ADAR1p150, we detected 2313 sites while ADAR1p110 was found to edit 2053 sites at frequencies ≥ 1% (Figure 2A, Figure 2 – table 2, Figure 2 – table 3). More than 40 % of the detected sites are efficiently edited by either isoform. Around one-third of editing sites is preferentially edited by ADAR1p150, and 25% of all sites are preferentially edited by ADAR1p110 (Figure 2B). Interestingly, ≈ 20 % seem exclusively edited by one or the other ADAR1 isoform (ADAR1p150: 346 and ADAR1p110: 257 ES) (Figure 2 - figure supplement 4). Still, despite the large overlap between ADAR1p150 and ADAR1p110 editing sites, the majority of the sites has a clear editing preference for one of the two isoforms (Figure 2C). Whilst ADAR1p110 is clearly responsible for the vast majority of intronic editing sites, the sites edited by ADAR1p150 show a more complex pattern. Interestingly, a large number of sites targeted by ADAR1 p150 are also located in intronic regions. This might be explained by the shuttling ability of ADAR1p150 or by redundant genomic annotations of intronic and exonic regions (Fritz et al., 2009). Still, ADAR1p150 is editing 3’UTRs with higher efficiency than ADAR1p110, consistent with its predominant cytoplasmic localization. Preferences for intronic regions and 3’ UTRs became even more prominent when only sites with a clear preference for editing by one of the two isoforms (sites that are edited 2,5 × better by one isoform than by another) were considered. There, ADAR1p110 edited almost exclusively intronic sites whereas 3’UTRs became a prominent target for ADAR1p150-editing (Figure 2D).

Next, we assessed editing in repetitive versus non-repetitive regions. ADAR1p150 edits more nonrepetitive parts (≈ 23%) in comparison to ADAR1p110 (≈ 16%) (Figure 2E). 3’UTRs that are preferentially edited by ADAR1p150 carry in general higher conservation than introns that are mostly edited by ADAR1p110. The vast majority of editing in repetitive regions by both ADAR1 isoforms occurs in short interspersed nuclear (SINE) elements (Figure 2F. Closer examination of editing events in repeats revealed also editing in LTR and long interspersed nuclear (LINE) elements (Figure 2F). Moreover, we studied ADAR1-editing in ‘hyperedited’ regions. Those regions contain many potential editing sites and frequently bear repetitive regions, in particular inverted SINEs. In our dataset, ADAR1p150 was a considerably greater editor of those hyperedited regions, which was becoming more evident when only sites with a clear preference for ADAR1p150 or ADAR1p110-editing (sites that are edited 2,5 × better by one isoform than another) were considered (Figure 2G). Most mRNA molecules spend the majority of their lifespan in the cytoplasm. Thus, ADAR1p150, which is mainly cytoplasmic, might have longer access to hyperedited regions than nuclear ADAR1p110.

Thus, our data show that ADAR1p110 edits mainly intronic repetitive regions, while ADAR1p150 is a powerful editor of 3’UTRs and hyperedited regions. ADAR1p150 is also more involved in editing of non-repetitive and thus potentially more conserved editing sites.

### ADAR-specificity is partially driven by its cellular localization

Having identified distinct editing patterns of ADAR1-isoforms, we wondered which features or characteristics of ADAR1 isoforms might drive the observed substrate specificities. ADAR1p150 differs from ADAR1p110 by an extended N-terminal region unique to p150. The p150-specific N-terminus harbors a nuclear export signal (NES) that determines the cytoplasmic localization and a Z-DNA binding domain (ZBD*α*) that can interact with double-stranded nucleic acids in Z-conformation. To evaluate if the cytoplasmic localization or the presence of ZBD*α* are driving ADAR1p150 specificity, we constructed mutant versions of both ADAR1 isoforms. Specifically, we generated nuclear ADAR1p150, cytoplasmic ADAR1p110 and ADAR1p150 bearing a P193A mutation. The P193A version was previously described in patients with Aicardi Goutière syndrome and is therefore believed to be an essential component of this motif (Rice et al., 2017). Furthermore, we constructed a cytoplasmic version of ADAR2 to fully evaluate the impact of cytoplasmic localization (Figure 3 - figure supplement 1). We confirmed the desired cellular localization of all mutants by immunofluorescent staining. ADAR1p150_P193A remained cytoplasmic as expected as the P193A mutation does not disturb NES function (Figure 3A). Next, we studied the impact of mislocalization on ADAR-editing specificity. Using Sanger sequencing, we evaluated editing in *Pum2* and *Deptor* upon transfection of different ADAR versions into MEFs^ADAR1-/- ;ADAR2 -/-^. *Pum2* and *Deptor* were edited by wt ADAR1p150 but not by wt ADAR1p110 and wt ADAR2. However, nuclear ADAR1p150 did not edit any of the selected sites, whereas cytoplasmic ADAR1p110 and even cytoplasmic ADAR2 did edit these substrates, as shown in the Sanger sequencing chromatograms. Surprisingly, the P193A mutation of ZBD*α* did not affect editing at both studied sites, which showed the same editing as wt ADAR1p150 (Figure 3B). This finding suggested that the specificity of ADAR1p150 isoform is prevalently driven by its cellular localization and, thus, might be substituted by any cytoplasmic editase.

**Figure 3:**
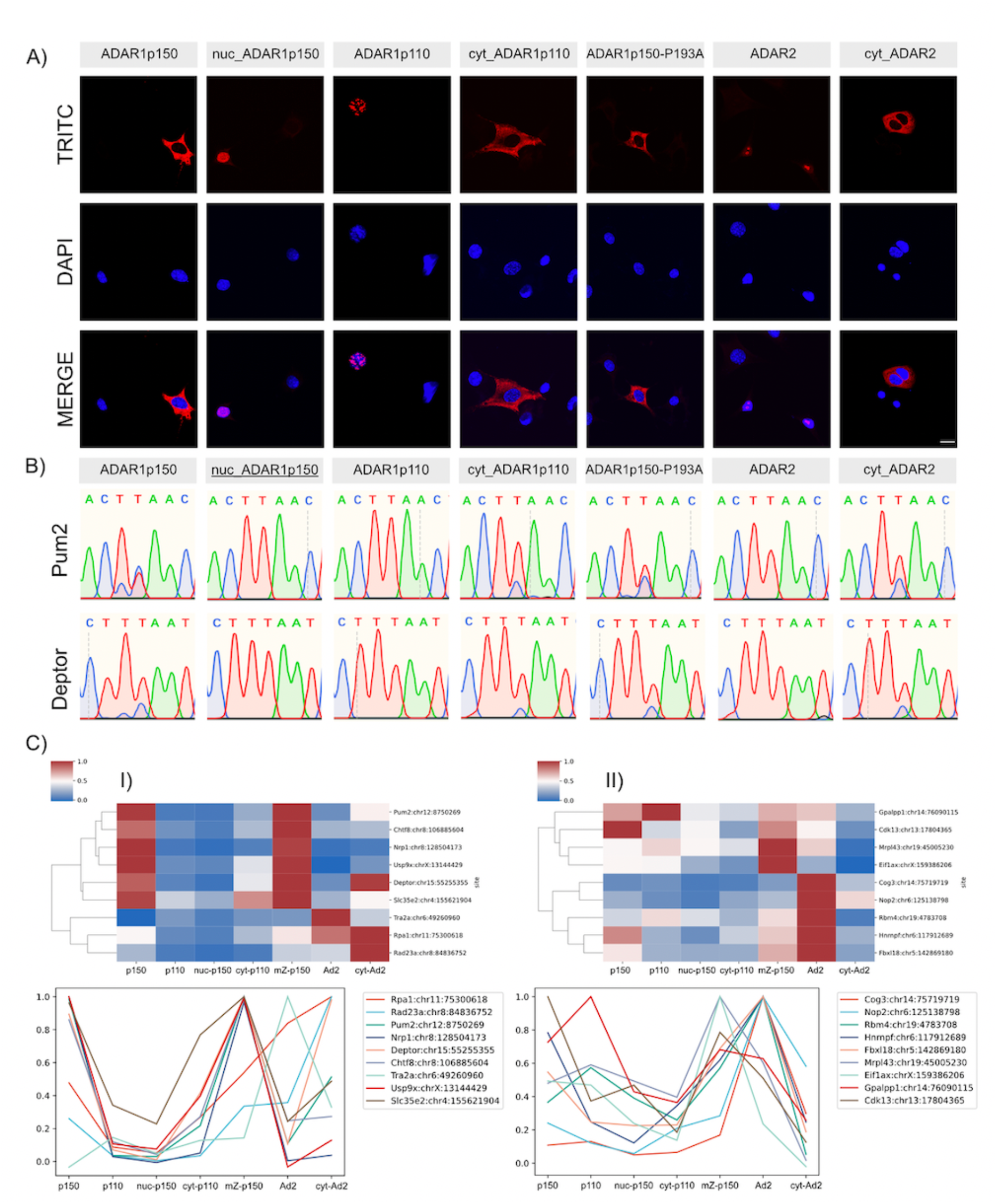
ADAR1-isoform specificity is partially driven by its cellular localization. A) Confocal microscopy images confirmed the expected cellular localization of mislocalized ADAR mutants. A P193A mutation in ZBDα does not affect the cellular localization, and ADAR1p150 P193A remains cytoplasmic. TRITC channel shows transfected constructs in confocal sections, and nuclear DNA is stained with DAPI. (Scale bar: 20 µm). B) Sanger sequencing traces evaluating the impact of mislocalization and ZBDα mutation on editing-specificity of ADAR1. Pum2 (chr12:8750269) and Deptor (chr15:55255355) are efficiently edited by ADAR1p150 but not by ADAR1p110. Cytoplasmic ADAR1p110 and cytoplasmic ADAR2 also edit both selected targets. In contrast, nuclear ADAR1p150 does not show any editing of selected targets. The P193A mutation does not affect editing of Pum2 and Deptor, leading to efficient editing in those substrates. C) Cluster maps and line plots of editing in selected editing sites detected by amplicon-seq. I) Editing sites identified as ADAR1p150 targets in MEFs. II) Editing sites identified as ADAR1p110 targets in MEFs. The majority of ADAR1p150 targets could be validated. The P193A mutation does not affect the editing pattern of the selected sites. The majority of the sites also shows a different level of sensitivity to cytoplasmic editing. Only limited editing by mislocalized cytoplasmic ADAR1p110 and cytoplasmic ADAR2 was seen in Nrp1 and Chtf8. In contrast, ADAR1p110 targets were barely verified with the exception of Gpalpp1. Instead, the majority of p110 targets was well edited by ADAR2. Surprisingly, many putative ADAR1p110 targets were also edited by ADAR1p150.

### Amplicon-seq of wt, mislocalized and mutated ADAR variants

To further assess the effect of mislocalized and mutated ADARs on their editing preferences, we performed amplicon-seq on a set of substrates (Figure 3 - figure supplement 2). To do so, we electroporated MEFs^ADAR1-/-; ADAR2-/-^ with all mutated and wt ADAR expressing vectors. An RFP expressing vector was used as a negative control. 17 editing targets were selected based on their preferential editing by ADAR isoforms and good expression, where 9 targets were preferentially edited by ADAR1p150 and 8 targets by ADAR1p110. Moreover, the chosen targets showed a highly diverse set of editing sites, including sites situated in exons, introns, 3’UTR, repeats and single-copy sequences, nonrepetitive regions (Figure 3 – table 1). Experiments were performed in triplicate using 5 µg of plasmid DNA for electroporation. A fourth replicate was included where 10 µg of plasmid DNA was electroporated. However, as this replicate did not contain the ADAR1P193A sample, it was not included in the final analysis. After STAR alignment, background signals occurring in the negative control were subtracted, and the remaining sites were subjected to further filtering. Sites edited to 0.5% by any mutant protein were collected into the final dataset. Overall editing in cells electroporated with 5 µg DNA was low. However, editing patterns remained similar as for cells transfected with 10 µg of DNA (Figure 3 - figure supplement 3, 4). Since absolute editing levels were heterogenous for the different constructs (Figure 3 - figure supplement 5), we normalized editing levels and set the maximum editing rate of a particular site to one (1) and calculated editing values for remaining editases proportionally (Figure 3C, Figure 3 – figure supplement 6) (Figure 3 -table 2). Using these normalized values, we could validate the majority of ADAR1p150-selected sites. However, individual sites in *Tra2a*, *Rpa1* and *Rad23* were preferentially edited by ADAR2 and cytoplasmic ADAR2. Our initial experiments did not include ADAR2, and hence, the occurrence of sites ADAR2-editing sites next to ADAR1p150 sites cannot be excluded. Nonetheless, the editing sites in *Rpa1* and *Rad23a* showed higher editing rates for ADAR1p150 than ADAR1p110 (Figure 3C). Quite surprisingly, the situation was less clear for ADAR1p110-targets. Selected sites of *Cdk13*, *Hnrnpf* and *Fbxl18* showed a clear preference for ADAR1p150, while the sites in *Gpalpp1* and *Eif1ax* showed a comparable preference for ADAR1p150 and ADAR1p110. Most interestingly, many sites that could be well-edited by ADAR1p110 were also highly edited by ADAR2, indicating preferential nuclear-editing of the selected sites (Figure 3C).

Surprisingly, the P193A mutation did not affect the ADAR1p150 editing pattern. The selected sites were edited to the same level as by wt ADAR1p150. Interestingly, the selected sites showed different preferences for cytoplasmic editing. While *Pum2*, *Deptor* and *Slc35e2* were edited by cytoplasmic ADAR1p110 and cytoplasmic ADAR2, *Chtf8* and especially *Nrp1* showed only limited editing by cytoplasmic ADAR1p110 and cytoplasmic ADAR2, although their natural editing is clearly mediated by ADAR1p150. Also, ADAR1p150 P193A edited *Nrp1* and *Chtf8* to a similar extend as wt ADAR1p150. The *Nrp1* editing pattern was also validated by Sanger sequencing (Figure 3 – figure supplement 7). Thus, *Nrp1* and *Chtf8* appear to be true ADAR1p150 editing substrates that specifically require ADAR1p150 for editing that cannot be substituted by other cytoplasmic editors.

To understand the characteristics of cytoplasmic and ADAR1p150 editing substrates, we analyzed their predicted folding using the Vienna RNA package and overlayed all detected sufficiently edited (≥0,5%) editing sites in the RNA structures (Figure 4, Figure 4 – figure supplement 1). Interestingly, even very closely spaced editing sites displayed highly variable editing preferences for wt and mutated ADARs.

**Figure 4:**
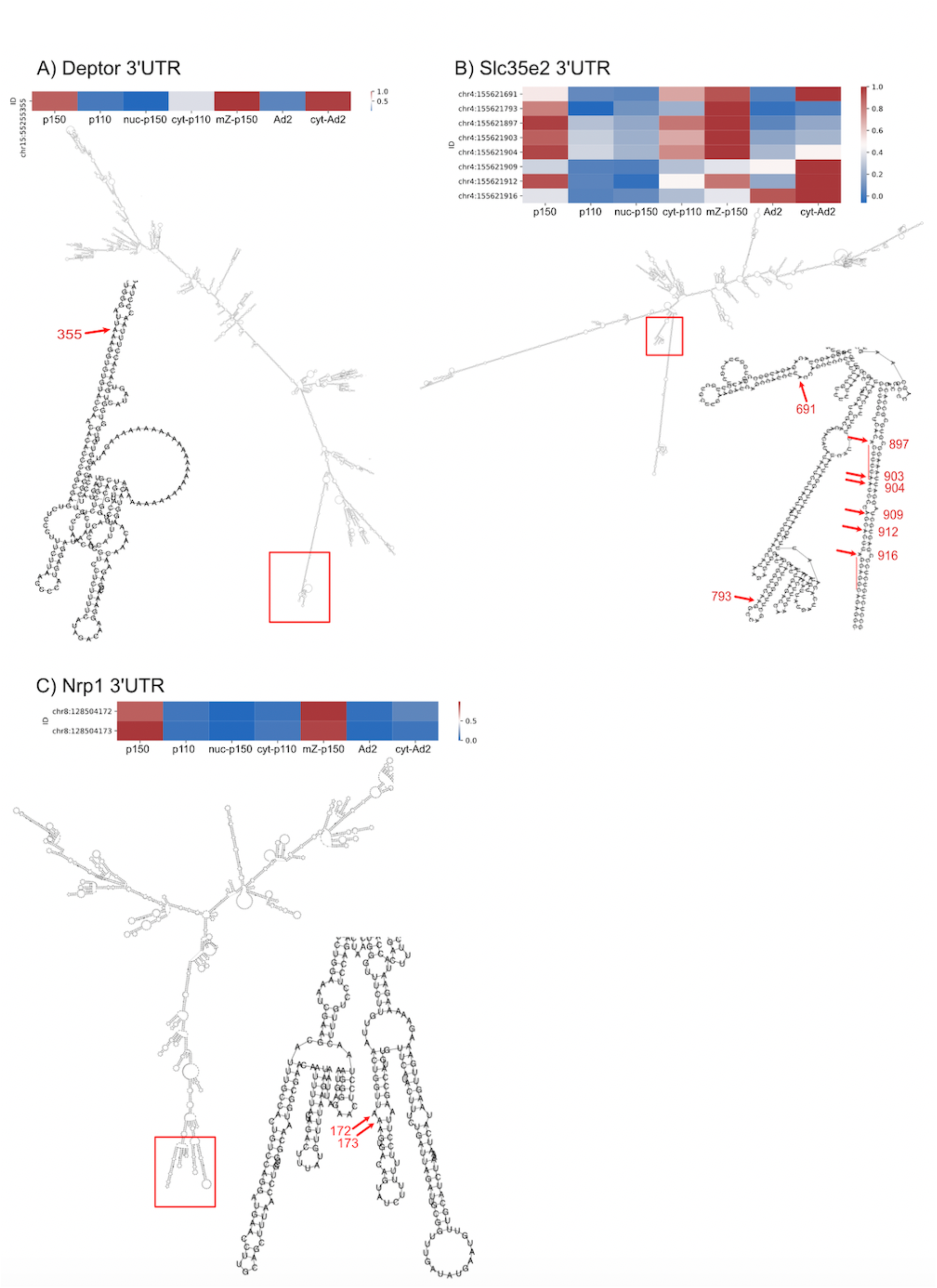
Heat maps of all editing sites detected by amplicon-seq in selected ADAR1p150 targets. Structure predictions were generated using Vienna RNA Package. Magnifications of edited regions display detected editing sites in the structure (labeled with the last 3 digits of the genomic coordinate).

Based on our observation, we speculated that substrates sensitive to general cytoplasmic editing (including cytoplasmic ADAR2) are embedded in long dsRNA stretches. This notion could be supported by the editing patterns seen in *Deptor*, *Pum2* and several sites of *Slc35e2* and *Rpa1* (Figure 4A, 4B, Figure 4 – figure supplement 1A, 1B). Authentic ADAR1p150 targets, on the other hand, seem to lie at the ends of long dsRNAs or within structures that are disrupted by bulges. Both *Nrp1* and *Chtf8* editing sites are located in more complex structures with many loops and bulges (Figure 4C, Figure 4 – figure supplement 1C)

Employing the amplicon-seq approach, we confirmed that ADAR1p150 specificity is indeed partially driven by its cytoplasmic localization. Based on this data, we suggest that mainly long-dsRNA structures are edited by any cytoplasmic editor. In contrast, substrates with complex structures are only edited by ADAR1p150, yet these sites are not affected by the P193A mutation in ZBD*α*.

### Mouse edited substrates and human MB-UV RIP-seq detect common substrates

To determine a possible overlap between mouse substrates edited by ADAR1 isoforms and human sequences bound by ADAR1 isoforms, we defined the overlap between both gene sets. We found an overlap of 445 genes for ADAR1p150 and 153 genes for ADAR1p110. This difference could be caused by the lower number of peaks detected by ADAR1p110 RIP-seq. Editing is very abundant in human and occurs in nearly all genes due to the high repeat content of the human transcriptome (Bazak *et al*., 2014a). Thus, we next focused on genes that were edited in non-repetitive regions. This resulted in 290 genes edited by ADAR1p150 and 142 genes edited by ADAR1p110. This way, we expected to find ADAR1 isoform-specific editing targets that are conserved between mouse and human. In doing so, we could identify 26 overlapping genes edited or bound by ADAR1p110. Strikingly, more than half (151) of the non-repetitive ADAR1p150 targets was common between human RIP-seq and mouse editing detection experiments (Figure 5, Figure 5 – table 1). These results are in agreement with our previous observation that ADAR1p150 is a powerful editor of 3’UTRs that are generally more conserved than introns which are predominantly edited by ADAR1p110.

**Figure 5:**
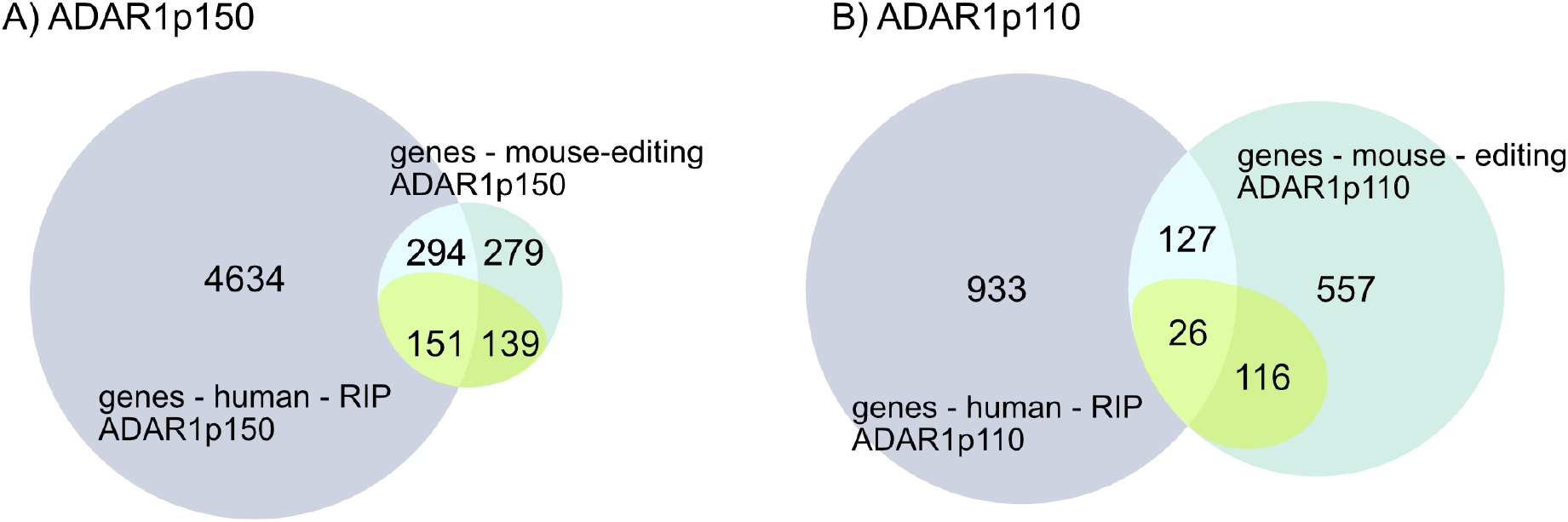
Intersection of regions identified in human ADAR1 RIP-seq with editing identified in mouse genes. A) More than half (151) of the genes containing non-repetitive editing sites were detected in both experiments performed with ADAR1p150. B) Only a limited overlap (26) was identified between two experimental approaches for ADAR1p110. In purple: genes that were identified by RIP-seq in human cells; turquoise: genes that were identified as editing targets in MEFs; yellow-green: genes containing editing sites in non-repetitive regions and those that were edited in MEFs.

## Discussion

In this study, we provide a robust overview of ADAR1-isoform-specific binding and editing characteristics. We identify RNAs that are selectively bound and edited by either ADAR1 isoform, while a large fraction of RNAs can be bound and edited by either isoform. Most interestingly, our study revealed that cellular localization is a major factor driving isoform specific editing patterns. Still, editing of isoform-specific substrates is not exclusively governed by the enzyme’s localization, indicating that also other factors, most likely RNA-binding or competing RNA-binding proteins, can trigger ADAR1 substrate-specificity.

Using a modified RNA-IP (RIP)-seq protocol that involved methylene blue and UV cross- linking, we could improve the pull-down efficiency of ADAR-associated sequences.

ADAR1 is a dsRNA binding protein that mainly interacts with the RNA backbone (Ryter and Schultz, 1998). Hence, conventional UV cross-linking is rather inefficient as this technique creates covalent bonds between a protein and free unpaired RNA-bases that are limited in dsRNA. We, therefore, applied methylene-blue mediated photo-crosslinking together with UV-crosslinking to achieve robust binding between ADAR1 and its RNA-substrates. Methylene blue intercalates into dsRNA, partially unwinds the dsRNA structure and thus, makes it accessible for photo-crosslinking (Liu *et al*., 1996). Moreover, additional UV-mediated cross- linking might further strengthen ADAR1-substrate binding and possible ssRNA-ADAR1 interactions (Wheeler et al., 2018). Using this technique, we found distinct binding regions for ADAR1-isoforms. While ADAR1p110 shows strongly enriched binding to introns, ADAR1p150 did efficiently capture exons and 3’UTRs.

We compared the outcome of our RIP-Seq results that were performed in HEK293 cells with the currently published ADAR1-isoform-specific editome that was also generated in HEK293 cells (Sun et al., 2021). To do so, we employed CrossMap (v0.5.4) (Zhao et al., 2014) to convert the set of identified editing sites to the GRCh38 assembly. Next, we defined testing groups: I) sites edited exclusively by ADAR1p150 (in at least 2 out of 3 replicas)- which yielded 2806 sites and II) sites edited by ADAR1p150 (in 2 out of 3 replicas) and ADAR1p110 (at least in 1 replica), which yielded in 4939 sites (Figure 6 – Table 1). Sun et al also identified a limited number of sites edited by ADAR1p110. However, since the editing specificity at these sites was not further confirmed we omitted them from further evaluation.

**Figure 6:**
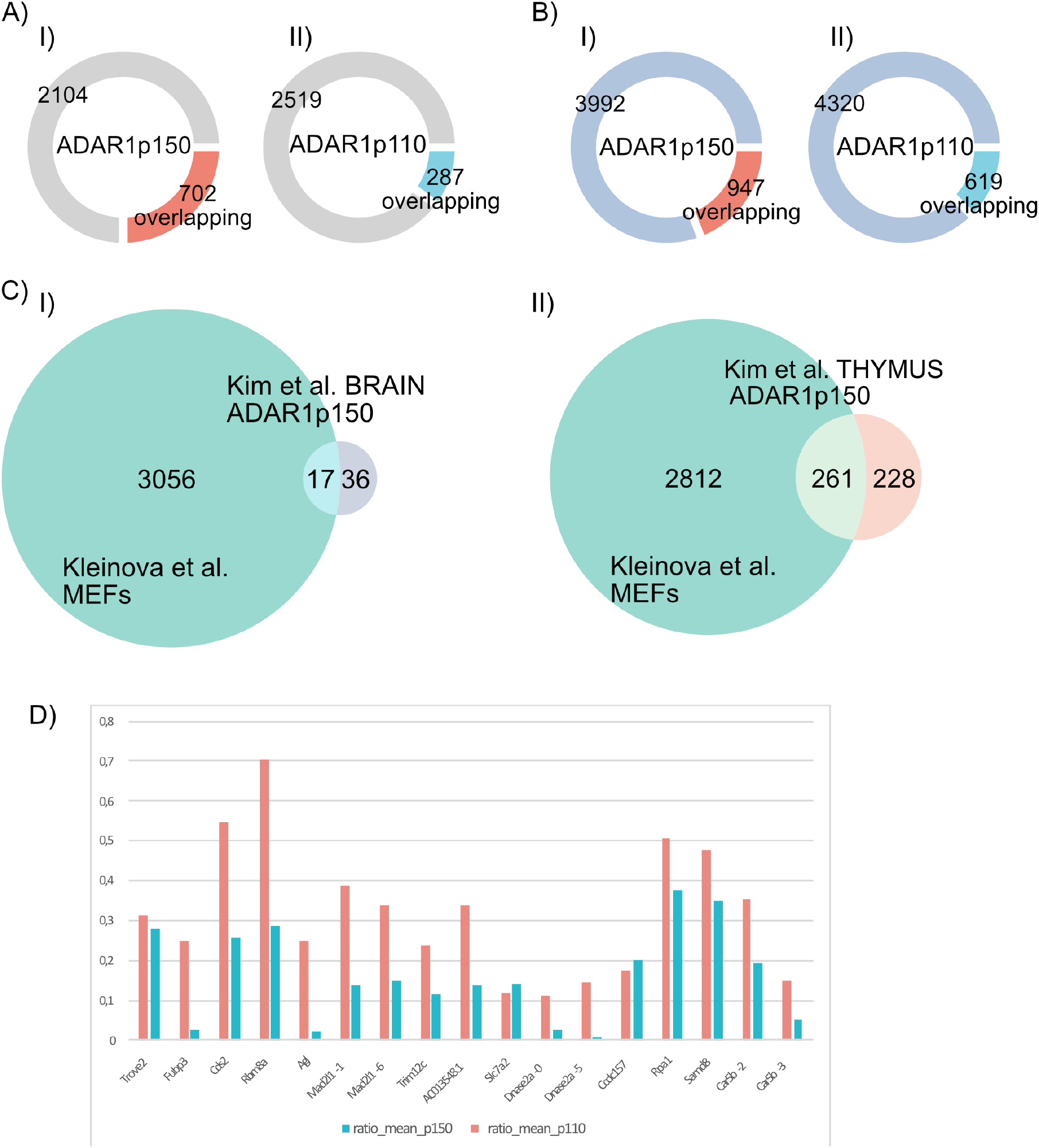
Discussion supporting figure. A), B) Intersection between the peaks identified in this study by MB-UV RIP-seq in HEK cells and editing sites identified by Sun et al. in HEK cells (Sun et al., 2021). Red and light-blue sections of doughnut plots display the portion of editing sites overlapping with RIP-seq peaks. A) I) Overlap of ADAR1p150-specific editing sites and ADAR1p150-peaks; II) Overlap of ADAR1p150-specific editing sites and ADAR1p110-peaks. B) I) Overlap of editing sites targeted by both ADAR1 isoforms and ADAR1p150-peaks; II) Overlap of sites edited by both ADAR1 isoforms and ADAR1p110-peaks. C) Comparison of ADAR1- editome identified in MEFs and ADAR1p150-specific editing sites detected in the brain and the thymus of ADAR1p110^-/-^ ADAR2^-/-^ . I) Editing sites identified as ADAR1p150-specific in the brain. II) Editing sites identified as ADAR1p150-specific in the thymus. D) ADAR1p150 and ADAR1p110-mediated editing ratios detected in MEFs for sites that overlap with ADAR1p150 sites in the brain identified by (Kim et al., 2021).

Next, using BEDTools-intersect (v2.30.0), we identified editing sites that directly overlap with RIP-seq peaks identified by us (Quinlan and Hall, 2010). Indeed, we found an overlap of ∼25% between sequence peaks defined by ADAR1p150 binding and editing sites that were selectively edited by ADAR1p150 as detected in the Sun et al., study (Sun *et al*., 2021) (Figure 6A). We also found an overlap of ∼19% between sequences bound by ADAR1p150 and editing sites targeted by both ADAR1p150 and p110. Consistently, the overlap between ADAR1p110- peaks and both examined editing site sets was considerable smaller: ∼10 % with ADAR1p150 editing sites and ∼ 12,5 % with the combined ADAR1p150 & p110 editing site set (Figure 6B) (Figure 6 – Table 2). Overall, the data show that ADAR1 p150 RIP-seq peaks have a larger overlap with sites edited by p150 while there seems less overlap with sites precipitated by ADAR1 p110 and those that are edited by ADAR1p150 and ADAR1p110. However, it should also be noted that RIP-seq identified significantly fewer peaks for ADAR1p110. This observation, together with preferential intronic binding of ADARp110 versus ADAR1p150 binding to exonic, 3’UTR regions, indicates that the identified peaks provide a valuable dataset of high specificity. Still, we cannot exclude that overexpression of ADAR1 leads to promiscuous binding and, hence ADAR1p110 could bind substrates that are otherwise exclusive for ADAR1p150. However, apart from the N-terminus of ADAR1p150, ADAR1- isoforms are identical (George and Samuel, 1999; Patterson and Samuel, 1995). Therefore, an overlapping affinity for the same substrates can be expected. RIP-seq of endogenous ADAR1 might improve the substrate-specificity of the procedure. Nonetheless, since endogenous ADAR1p150 is expressed at a very low level unless the cells are stimulated with IFN (Patterson and Samuel, 1995) and specific antibodies recognizing only the N-terminus of ADAR1p150 are scarce. Therefore, experimental conditions would require adequate optimization to specifically detect binding sites of ADAR1 isoforms.

After having established ADAR1 isoform-specific binding substrates in human cells we also examined ADAR1-isoform specificity in the mouse. Having the advantage of ADAR deficient mouse cells as a tool, we restored ADAR1 expression in those cells. Here the ADAR1- isoform-specific editome was determined to gain insight into isoform-specific targets.

More than 100,000 known editing sites identified mostly in mouse brain are currently available on REDIportal (http://srv00.recas.ba.infn.it/atlas/search.html) (Picardi *et al*., 2017).

Of these, we detected 9,400 sites by restoring ADAR1-mediated editing in ADAR-deficient MEFs. Using additional stringent filtering criteria, we obtained a comprehensive, high-quality dataset of more than 3000 editing sites. We found ∼2000 editing sites efficiently edited by one or another ADAR1-isoform. While ADAR1p110 edits almost exclusively in introns, ADAR1p150 seems a powerful editor of 3’UTRs. As expected, the majority of editing sites is located in SINE elements. Nevertheless, ADAR1p150 showed a bigger portion of editing in non-repetitive regions, which suggests that ADAR1p150-mediated editing might be more conserved among species. Furthermore, ADAR1p150 is also more involved in the editing of hyperedited regions, most probably due to the spatio-temporal advantage of its cytoplasmic localization.

Although we identified ADAR1p110 editing sites, we cannot fully exclude that these sites might also be edited by ADAR1p150. The RNA-seq depth is, in general, lower in intronic regions that are preferentially edited by ADAR1p110 sites therefore, deeper sequencing might reveal editing mediated also by ADAR1p150. Indeed, the editing sites in *Gpalpp1* (chr14:76090115) that were identified to be exclusively edited by ADAR1p110 could subsequently be shown to be edited by ADAR1p150 as well. However, this required amplicon-seq with several times higher coverage (Figure 3C, Figure 3 – Table 1). A similar observation was made by Sun et al. when studying editing in HEK cells. Although several putative ADAR1p110 sites could be identified initially, subsequent amplicon sequencing revealed that many sites can also be targeted by ADAR1p150 (Sun *et al*., 2021). Thus, ultra-deep sequencing would be needed to identify true ADAR1p110 targets.

ADAR1 deficient mice and loss of function mutations in human ADAR1 share similar phenotypic characteristics, including massive IFN-overproduction (Mannion *et al*., 2014). Mouse-knockout studies showed that specific editing activity of ADAR1p150 is essential to prevent undesired endogenous RNA-sensing by MDA5 (Liddicoat *et al*., 2015; Pestal *et al*., 2015). Therefore, similar characteristics must be common for human and mouse ADAR1p150- mediated editing. In our current study, we employed two different approaches: I) ADAR1-RIP-seq performed in human cells and II) isoform-specific reconstitution of ADAR1 editing in mouse MEF cells. To further compare these two approaches, we focused on genes with editing sites in non-repetitive regions, assuming that such sites may exhibit more genomic conservation. While only ∼18 % of such genes overlap between the two experimental strategies for ADAR1p110, more than half of such genes are common between ADAR1p150 RIP-seq and ADAR1p150 editing analysis. Closer inspection revealed many interesting ADAR1p150 substrates, including a highly-conserved editing site in *Azin1* (chr15:38491612) recently described as ADAR1p150 substrate also by Kim et al. (Figure 6 – figure supplement 1A) (Kim *et al*., 2021). Editing in *Azin1* is extensively studied for its role in many cancer types and lately also for its involvement in hematopoietic stem cell differentiation (Hu et al., 2017; Okugawa et al., 2018; Shigeyasu et al., 2018; Wang et al., 2021).

Still, it appears unlikely that one individual substrate is responsible for the phenotype of ADAR1 deficiency. Instead, more general editing characteristics might cause the indispensability of ADAR1p150-editing. We could clearly observe a correlation between ADAR1-editing in mouse and ADAR1-binding distribution in human cells. While ADAR1p110— editing and binding fall mostly into intronic regions, 3’UTRs seem a common prominent substrate for ADAR1p150 binding and editing. 3’UTRs often contain inverted SINE elements and thus form dsRNA structures that present ideal substrates for ADAR1 with many putative editing sites (Bazak *et al*., 2014a; Faulkner et al., 2009). Importantly, such long dsRNA stretches also present ideal triggers for MDA5-activation (Ahmad et al., 2018). In our TRIBE experiment, these hyperedited regions were preferentially edited by ADAR1p150. This indeed raises the question of whether sufficient editing of these hyperedited 3’UTRs is a key and essential feature of ADAR1p150-mediated editing. A recent study from Kim et al. showed only limited editing sites that are exclusively edited by ADAR1p150 to prevent MDA5-activation (Kim *et al*., 2021). In this study, mouse knockout models including an ADARp110 specific knockout in the presence or absence of an additional ADAR2 deficiency were used to characterize the remaining ADAR1p150-mediated editing in brain and thymus. We intersected the exclusive-ADAR1p150 editing pattern identified in that study with the ADAR1 p150 editome identified here in MEFs. We found a big overlap between the datasets. 23 sites, out of 53 in the brain and 312 out of 489 in the thymus (Figure 6C). Although all sites originally identified in the brain showed a great preference for ADAR1p150-editing in MEFs, none of them was exclusively edited by p150 in our setup (Figure 6D). In the thymus editing set, we found 23 sites exclusive for ADAR1p150 editing and additional 22 sites with a high (more than 10fold change) preference for ADAR1p150-editing in isoform-transfected MEFs (Figure 6 – Supplementary figure 2; Figure 6 – Table 3). We hypothesized that the limited number of exclusive ADAR1p150 sites identified by Kim et al. might result from a limited sequencing depth and might be increased by deeper sequencing. In fact, the coverage of examined sites was two times lower than in our experiment, although the total number of sequenced reads was comparable (Figure 6 – figure supplement 3). Most overlapping sites fall into hyperedited 3’UTRs, which might again indicate that those are the essential ADAR1p150 sites that have to be masked by editing to prevent MDA5-triggering under physiological conditions.

However, the main driver of ADAR1p150-specificity still remained elusive. Predominant cytoplasmic localization and the ZBD*α* are two exclusive features of ADAR1p150. Nevertheless, their involvement in editing or binding-specificity is not understood. Several mutations of ZBD*α*, mainly P193A, were connected to AGS (Rice *et al*., 2017). ZBD*α* recognizes the unusual Z-conformation in RNA or DNA that is promoted mainly by CG repeats. Therefore ZBD*α* might contribute to the selectivity of ADAR1p150 (Herbert and Rich, 1999; Koeris et al., 2005; Schwartz et al., 1999). Moreover, several recent studies that introduce mutations in ZBD*α* in mice show that ZBD*α* is crucial to suppress MDA5 activation and contributes to editing selectivity of ADAR1p150 (de Reuver *et al*., 2021; Maurano et al., 2021a; Nakahama *et al*., 2021; Tang *et al*., 2021). A mutation of ZBD*α* mainly affects editing in SINE elements (de Reuver *et al*., 2021) which is in accordance that inverted SINEs form long dsRNA with putative Z-RNA conformation (Herbert, 2019). While ZBD*α* is intensively studied, the role of typical cytoplasmic localization of ADAR1p150 on its editing specificity remains unclear. Nuclear editing occurs co-transcriptionally, therefore only limited time is available for editing of substate-sites (Bentley, 2014; Licht *et al*., 2016). In contrast, most mRNAs spend most of their lifespan in the cytoplasm. Hence cytoplasmic ADAR1p150 might have the spatio- temporal advantage to edit hyperedited regions. Interestingly, we observed that many ADAR1p150-specific sites are edited by mutant versions of ADAR1p110 and ADAR2 that localize to the cytoplasm. This finding suggests that the specificity of ADAR1p150 is at least partly driven by its cytoplasmic localization. However, a few sites, especially sites in *Nrp1*, seem insensitive to cytoplasmic versions of other editing enzymes and require exclusive ADAR1p150-editing. Based on RNA-folding predictions, it appears that substrates sensitive to cytoplasmic editing form extended dsRNA structures – an ideal ADAR-substrates. In contrast, *Nrp1* editing sites specific for ADAR1p150 are localized in more complex structures (Figure 4C).

*Nrp1* is a cell surface glycoprotein that serves as a receptor for VEGF however, recent studies showed that *Nrp1* is also SARS-CoV-2 coreceptor that facilitates virus cell entry and infectivity (Cantuti-Castelvetri et al., 2020; Hu et al., 2021; Soker et al., 1998). The role of editing in 3’UTR of *Nrp1* has not been studied even though the editing sites and surrounding region share great conservation amongst mammals (Figure 6 – figure supplement 1B). While editing of the 3’UTR (chr10: 128504171, 128504172, 128504173) in mouse is annotated in REDIportal, the corresponding human sites are not included in REDIportal or DARNED. Nonetheless, the corresponding human site for ch10:128504173 - chr10:33467752 (hg19) was identified as an exclusive ADAR1p150 target in HEKs by Sun et al., (Sun *et al*., 2021). This indicates that the *Nrp1* 3’UTR carries conserved editing sites that are edited by ADAR1p150. Additional studies would be required to assess the possible effect of 3’UTR editing on *Nrp1*- biology, stability, expression and miRNA binding pattern. Apart from *Nrp1*, an examined site in *Chtf8* seems to be also insensitive to nonspecific cytoplasmic editing and similar to *Nrp1* it is located in a region predicted to fold into a rather complex structure. Our observation is based on the amplicon-seq experiment that provides enormous sequencing depth at limited sites. Deep RNA-seq would be required to detect more ADAR1p150 specific sites that are otherwise insensitive to cytoplasmic editing to obtain a comprehensive picture of ADAR1p150 selectivity.

Surprisingly, a P193A mutation of ZBD*α* in ADAR1p150 only marginally affected editing of the studied substrates. A recent study by Maurano et al. showed that P195A, mimicking human P193A is not mutagenic on its own but is haploinsufficient (Maurano *et al*., 2021b). Similarly, mice bearing homozygous N175A/Y179A (mimicking human N173A/Y177A), which considerably disturbs Z-RNA binding (Placido et al., 2007; Schade et al., 1999), are viable. Nonetheless, ADAR1^N175A/Y179A^ triggers an MDA5/MAVS-related IFN-response and shows reduced editing of SINEs (de Reuver *et al*., 2021; Tang *et al*., 2021). Consistently, human N173A that is homologous to mouse N175 was already identified in an AGS patient, a mutation corresponding to Y179 has not been detected in humans (Rice *et al*., 2017). AGS patients with a P193A mutation in the compound heterozygous state with a more severe dysfunctional ADAR1 allele have also been identified (Rice *et al*., 2012; Rice *et al*., 2017). In conclusion, ZBD*α* probably facilitates editing of certain substrates. However, the pathologic effects of mutations in ZBD*α* might be overcome by higher expression of cytoplasmic editases that substitute for disrupted binding by mutated ZBD*α*. ADAR1p150 is interferon-inducible, thus, once activated, ADAR1p150 is produced in high amounts which then leads to a great amount of cytoplasmic editase that massively edits substrates in the cytoplasm. This might explain the lack of phenotype in ADAR1^P195A/ P195A^ mice and the lack of AGS patients carrying homozygous P193A mutations (Maurano *et al*., 2021b; Rice *et al*., 2017).

In our study, we determined the characteristic of ADAR1-isoform-specific editing and binding. We obtained a comprehensive, high-quality dataset that identifies specific and overlapping substrates between ADAR1p150 and p110. Our data indicate that editing specificity of ADAR1p150 is at least partly driven by its cytoplasmic localization. A proper genetic mouse model with a physiological expression of either cytoplasmic ADAR1p110 or cytoplasmic ADAR2 in an otherwise editing-free background would be required to fully estimate the role of the cytoplasmic localization and ZBD*α* in ADAR1p150-specificity. Additionally, we identified sites with no obvious sensitivity to unspecific cytoplasmic editing; putative exclusive ADAR1p150 sites. Based on our data, we hypothesize that promiscuous cytoplasmic editing occurs mostly in long dsRNA stretches, whereas exclusive ADAR1p150 editing sites lie in structures with more complex folding patterns. Loss of ADAR1 has recently been shown to help to overcome PD1 checkpoint blockade (Ishizuka et al., 2018). It will thus be interesting to see whether the substrates identified here are involved in overcoming PD1 blockage when expressed in an unedited state.

## Methods

### Cell culture and MEF generation

Human embryonic kidney 293 cells (HEK 293) were maintained in high-glucose Dulbecco’s Modified Eagle Medium (DMEM) (Thermo Fisher Scientific, Waltham, Massachusetts) supplied with 10% fetal bovine serum, pyruvate and L-glutamine.

#### Generation and culture of Mouse embryonic fibroblasts (MEFs)

Mice carrying a homozygous deletion of *Adarb1* (Adarb1^-/-^) that were rescued by a pre-edited version of glutamate receptor subunit 2 *Gria2*^R/R^ and carrying a simultaneous heterozygous deletion of *Adar (Adar^+/-^)* were bred. After 11 days, pregnant females were sacrificed, and embryos of genotype *Adar^-/-^; Adarb1^-/-^*; *Gria2^R/R^* were macerated and cultured in DMEM supplemented with 20% FBS. Early passage MEFs were immortalized using lentiviral transduction (Zhao et al., 2010).

### RNA extraction

RNA was extracted using TriFAST^TM^ (VWR, Peqlab, Radnor, PA, USA) according to the manufacturer’s suggestions. Isolated RNA was treated with DNaseI (New England Biolabs, Ipswich, Massachusetts) and subsequently purified by phenol: chloroform, chloroform extraction and precipitated with ethanol.

### ADAR-RIP

HEK cells were seeded at a density of 6,5 × 10^6^ cells per 150mm dish. 24 hours later, 2 dishes each were transfected per condition with 38 µg of plasmid DNA using 90 µl of linear polyethyleneimine (PEI) (Polysciences Warrington, PA, USA) and incubated for 24 hours. Dishes were kept on ice and washed twice with ice-cold 1×PBS. Cells were subjected to either native RIP or cross-linked RIP procedure:

*Native ADAR RIP* was performed as described previously (Hsieh et al., 2014). Only one 100mm dish per condition was subjected to the protocol (transfection mixture was scaled down proportionally). In short, cells were scraped and lysed in 1 ml of ice-cold polysomal lysis buffer (PLB) (100 mM KCl, 5 mM MgCl2, 10 mM HEPES (pH 7.0), 0.5% NP40, 1 mM DTT, 50 U of RNase inhibitor (New England Biolabs, Ipswich, Massachusetts), protease inhibitor cocktail (Complete Mini, Roche, Merck, Kenilworth, NJ, USA)). The cell suspension was passed 8 times through a 27,5-G needle. The cell lysate was spun for 15 min, 16 000g at 4 °C. The supernatant was precleaned with 40 µl pre-washed (2 × with PLB) Dynabeads® Protein A (Thermo Fisher Scientific, Waltham, Massachusetts) for 1 hr at 4 °C. The input sample was collected. anti-FLAG antibody (F7425, Sigma Aldrich, Merck, Kenilworth, NJ, USA) was added to each sample and rotated overnight at 4 °C. 30 µl of Dynabeads® Protein A (pre-washed 2 × in PLB) were added to each sample and rotated for another hour at 4 °C. Beads were subsequently washed 3 times with PLB. 10 × DNaseI reaction buffer was added, and samples were treated with 10 µl of DNase I (New England Biolabs, Ipswich, Massachusetts) for 15 min at 37 °C. RNA was extracted with 1 ml of TRIFAST as described above for general RNA extraction.

#### Cross-linked ADAR1 RIP

For formaldehyde cross-linking, 0,1 % formaldehyde in 1 × PBS was added to cells and incubated for 10 minutes at room temperature with gentle mixing. Cross-linking was quenched by adding one-tenth of quenching buffer (2.5 M glycine and 25 mM Tris) (Ricci *et al*., 2014).

Methylene Blue cross-linking was performed by adding 18 ml of 3µg/ml Methylene Blue in 1 × PBS. Cells were kept on ice and subsequently exposed to visible light for 30 minutes (Kaiser Prolite 5000). Ultraviolet cross-link 2 × 800 mJ was additionally applied (UVP Crosslinker, Analytic Jena GmbH, Jena, Germany).

Cross-linked cell pellets were kept at -80 °C until cell lysis. The IP-protocol was modified from Ricci *et al*. (Ricci *et al*., 2014). In short, cell pellets were lysed in 1.2 ml (per 2×150 mm dishes) of hypotonic lysis buffer (20 mM Tris-HCl, pH 7.5, 15 mM NaCl, 10 mM EDTA, 0.5% NP-40, 0.1% Triton X-100, 0.1% SDS and 0.1% sodium deoxycholate, protease inhibitor cocktail (Complete Mini, Roche, Merck, Kenilworth, NJ, USA) and 160 U of RNase inhibitor (New England Biolabs, Ipswich, Massachusetts)) on ice for 10 minutes.

The suspension was sonicated (Sonopuls HD 2070, Bandelin, Berlin, Germany) at 40% amplitude for a total of 60 s, 2×30 s on ice with a 20 s break and subsequently treated with 8 µl/sample Turbo DNaseI for 10 min at 37 °C (Invitrogen^TM^, Thermo Fisher Scientific, Waltham, Massachusetts). NaCl was adjusted to 150 mM, and the lysate was cleared by two centrifugation steps at 15.000 g for 10 minutes at 4°C. The input samples were collected. The lysate was incubated for 2 hours at 4°C with 180 µl of anti-flag agarose beads (50 % slurry, ANTI-FLAG® M2 Affinity Gel, Merck, Kenilworth, NJ, USA), prewashed 3 × with 1 ml isotonic wash buffer (IsoWB) (20 mM Tris-HCl, pH 7.5, 150 mM NaCl, and 0.1% NP-40). Next, beads were washed 2 × with 1 ml IsoWB + 0.1% SDS and 0.1% sodium deoxycholate, then 2 × with 1 ml with IsoWB. Protein-RNA complexes were eluted with 300 µl of PK-7M Urea buffer (200 mM Tris-HCl pH 7.4, 100 mM NaCl, 20 mM EDTA, 2% SDS, and 7 M Urea) at 25°C for 2 hr with 1000 rpm. Proteins were digested with 2mg/ml of preincubated proteinase K for 2 hours, at 25 °C with 1000 rpm agitation. RNA was extracted with TRIFAST, treated with DNaseI, followed by Phenol-Chloroform extraction. 10 ng of MB+UV-RIP- and corresponding input- RNA was subjected to rRNA-depletion with NEBNext^®^ rRNA Depletion Kit (Human/Mouse/Rat). cDNA libraries were synthesized using NEBNext^®^ Ultra^™^ II Directional RNA Library Prep Kit for Illumina^®^ (Both: New England Biolabs, Ipswich, Massachusetts) and subsequently sequenced in a paired-end mode with 75bp read length on NextSeq500 (Illumina, San Diego, CA, USA) (to obtain ≈ 10-15 mil mappable reads per RIP and input sample).

### RT-qPCR for ADAR-RIP evaluation

20 ng of RIP-RNA and input-RNA was spiked in with 25 ng of fly RNA and reverse transcribed using Maxima H MinusTranscriptase (Thermo Fisher Scientific, Waltham, Massachusetts) and random hexamer primers. qRT-PCR was subsequently performed with primers specific for edited genes and fly sequences using Luna Universal qPCR Mix (New England Biolabs, Ipswich, Massachusetts) and Biorad CFX Connect^TM^ Real-Time PCR Detection System (BioRad, Hercules, CA, USA). qPCR results were normalized on fly-spiked in RNA and a relevant input sample.

### ADAR-RIP NGS data analysis

Sequenced reads were aligned to the human reference genome (H.sapiens/GRCh38) using STAR aligner version 2.5.2a (Dobin et al., 2013). RIP-seq peak calling was performed with Genrich version 0.6 (https://github.com/jsh58/Genrich, parameters: -q 0.05 -a 200). Genrich was originally designed for genomic enrichment assay. The peaks were called on all replicates collectively for each ADAR1p150, ADAR1p110 and mock transfection. Respective input RNA samples were used as control samples. To allow strand-specific peak calling by Genrich, forward and reverse strands of a chromosome were treated separately.

Analysis was performed on peaks found in two or more replicas where p-values were below 0.1. Peaks appearing in empty-transfection control (PEI) were extracted from the final peak- set using bedtools (version 2.29.2).

### Restoring ADAR1 expression in editing-deficient cells

0,7 × 10^6^ MEFs were electroporated with 5 µg plasmid DNA using a Neon Transfection System (Thermo Fisher Scientific, Waltham, Massachusetts) with parameters: 1350 V, 30 mS, 1 pulse and 100 µl Neon tip. Cells were harvested after 24 hours. 100 ng of DNaseI-treated RNA was rRNA depleted using NEBNext^®^ rRNA Depletion Kit (Human/Mouse/Rat). cDNA libraries were generated with NEBNext^®^ Ultra^™^ II Directional RNA Library Prep Kit for Illumina^®^ (New England Biolabs, Ipswich, Massachusetts) and subsequently sequenced in paired-end mode with 150- bp read length on a NextSeq500 to obtain ∼40 mio reads (Illumina, San Diego, CA, USA).

### NGS data processing and A-to-I editing detection: REDItools

Sequenced reads were inspected with FASTQC (http://www.bioinformatics.babraham.ac.uk/projects/fastqc) and cleaned using FASTP to remove adaptors as well as low-quality regions (Chen et al., 2018). Cleaned reads were mapped onto the mouse reference genome (GRCm38 assembly), using STAR aligner (Dobin *et al*., 2013) and a list of known splice junctions from Gencode. Uniquely mapped reads in BAM format and a list of known mouse RNA editing sites from REDIportal were then passed to REDItools to profile known RNA variants per sample (Picardi *et al*., 2017; Picardi and Pesole, 2013).

### Construction of mislocalized ADAR vectors and immunostaining

Gibson assembly (NEBuider Hifi DNA Assembly Cloning kit, New England Biolabs, Ipswich, Massachusetts) and mutagenesis primers were used to generate mutated flag-ADAR mammalian expression vectors. In short, nuclear ADAR1p150 was prepared by deletion of full NES (ID: P55265-1, 125-150: CLSSHFQELSIYQDQEQRILKFLEEL) (Strehblow *et al*., 2002). The nuclear localization of the mutant was additionally supported by inserting a strong SV-40 NLS (PKKKRKVEDP) placed instead of deleted NES. ADAR1p150 ZBD*α* mutant was made by substitution of P193 to A. Cytoplasmic ADAR1p110 was prepared by introducing a single mutation at the position R801A – the crucial residue of ADAR1-NLS (Barraud *et al*., 2014). Cytoplasmic ADAR2 was produced by deletion of its N-terminal part bearing NLS (ID: P51400- 1, 1-72) (Behm et al., 2017; Desterro et al., 2003) and the cytoplasmic localization was additionally supported by placing minimal ADAR1p150 NES (CLSSHFQELSIY, 125-136) (Poulsen *et al*., 2001) instead of the deleted NLS.

Localization of resulting plasmids was assessed with immunofluorescence staining using an anti-flag antibody (F7425, Sigma Aldrich, Merck, Kenilworth, NJ, USA) in combination with Alexa 546 secondary antibody (Invitrogen, Thermo Fisher Scientific, Waltham, Massachusetts) and counterstained with DAPI. Microscopic confocal sections were taken on Confocal Laser Scanning Microscope FV3000 (Olympus, Tokyo, Japan) and images were processed with ImageJ.

### RT-PCR and A-to-I editing evaluation using Sanger sequencing

DNaseI-treated RNA was reverse transcribed using M-MuLV reverse transcriptase (New England Biolabs, Ipswich, Massachusetts) and random hexamer primers. cDNA fragments spanning selected editing sites in *Azin1, Pum2, Deptor* and *Nrp1* were amplified (see table: primers), gel eluted, and sent for Sanger sequencing (Eurofins, Luxembourg). The chromatograms were evaluated with SnapGene Viewer software.

### Amplicon sequencing and data analysis

MEFs were electroporated with indicated plasmids for 24 hours. After RNA extraction and cDNA, synthesis amplicons were generated with OneTaq or Q5 Hifi DNA polymerase (both: New England Biolabs, Ipswich, Massachusetts) by 2-step PCR. First, cDNA fragments were amplified with gene-specific primers containing a part of the Illumina adaptor sequence. Second, fragments were barcoded by PCR with primers containing the adapter sequence and unique indexes for multiplexing. Amplicons were gel-purified, pooled and sequenced in paired-end mode with 150 bp read-length on NextSeq550 (Illumina, San Diego, CA, USA).

Reads were adapter-clipped using Cutadapt (Martin, 2011) and aligned to the mouse genome mm10 with STAR using public server at usegalaxy.org (Afgan et al., 2018; Dobin et al., 2012). A-to-I transitions were then detected and quantified with Pysam (version 0.16.0.1) (Li et al., 2009) *<*http://www.ncbi.nlm.nih.gov/pubmed/19505943) employing REDIportal set of mouse known-editing sites (http://srv00.recas.ba.infn.it/atlas/index.html) (Picardi *et al*., 2017).

### Structures of RNA

RNA-structure was predicted using RNAfold 2.0 with with default parameters via the Vienna RNA websuite (Gruber et al., 2008; Lorenz et al., 2011).

### Data deposition

NGS data generated in this study are available at GEO with the following accession numbers: ADAR1 RIP-seq data: GSE188937 Editing data of transfected MEFs: GSE188842

## Acknowledgement

The authors would like to thank Katarina Milanovic for her help with amplicon-seq. Tanja Rohr and Alwine Hildebrandt for excellent technical support to. We also thank Cornelia Vesely, Mamta Jain, Andrea Tanzer and Konstantin Licht for helpful discussions and valuable input.

The ADAR1 and ADAR2 vectors were a kind gift of Mary Anne O’Connel (Ceitec, Brno, Czech Republic) and Ron Emeson (Vanderbilt University, Nashville, TE).

Illumina sequencing was performed at the Core Facilities of the Medical University of Vienna, a member of VLSI. The Amplicon-seq experiment was performed by the Next Generation Sequencing Facility at Vienna BioCenter Core Facilities (VBCF), member of the Vienna BioCenter (VBC), Austria.

## Supplementary figures

**Figure 1 – figure supplement 1.**
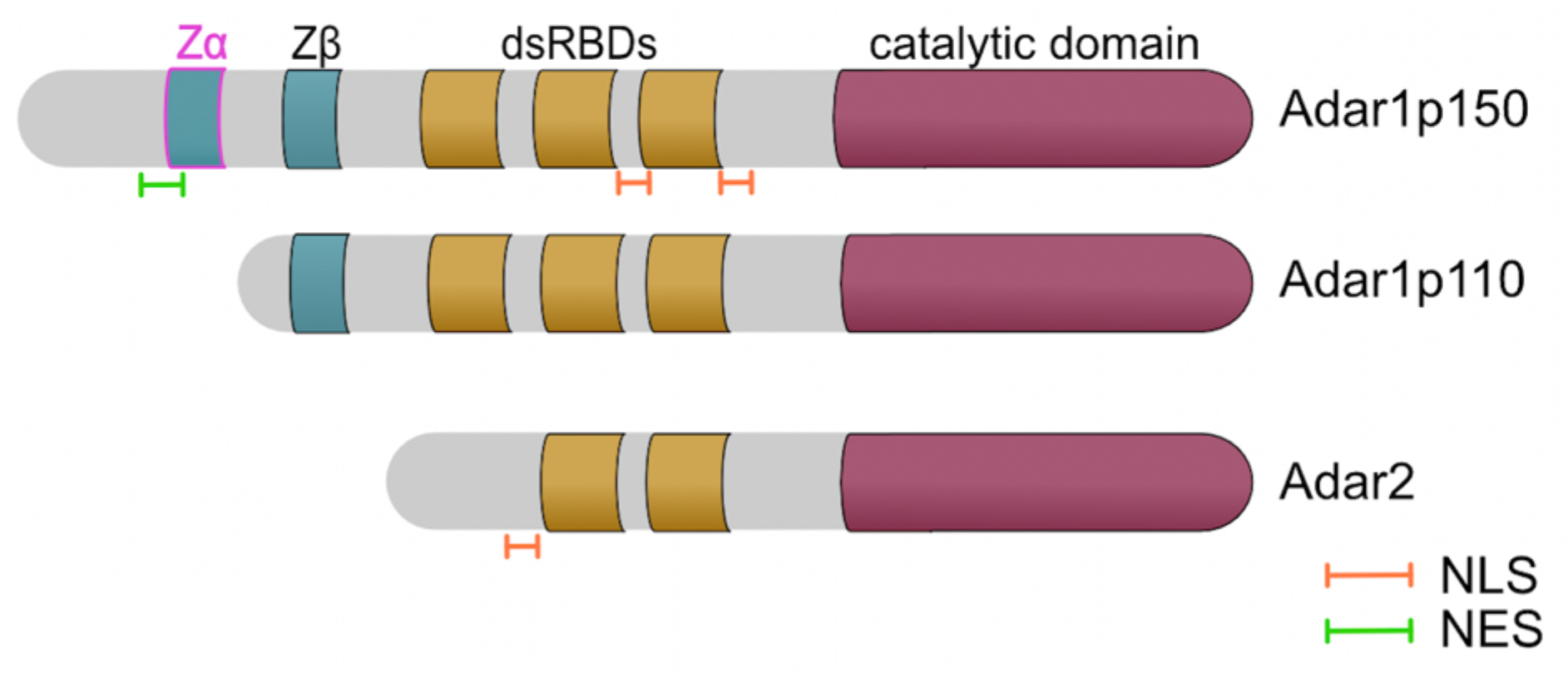
Domain organisation of active mammalian ADAR proteins: ADARs contain a conserved C- terminal catalytic-deaminase domain and two or three double-stranded RNA binding domains (dsRBD). Whilst both ADAR1 isoforms have Z-DNA binding domain *β* (Z*β*), full-length ADAR1p150 isoform also bears N-terminal Z-DNA binding domain *α* (Z*α*) and a nuclear export signal (NES). All ADAR variants possess nuclear localisation signals (NLS). While the NLS of ADAR2 is localised at the N-terminus, the NLS of ADAR1 is bimodular assembled around the third dsRBD.

**Figure 1 – figure supplement 2.**
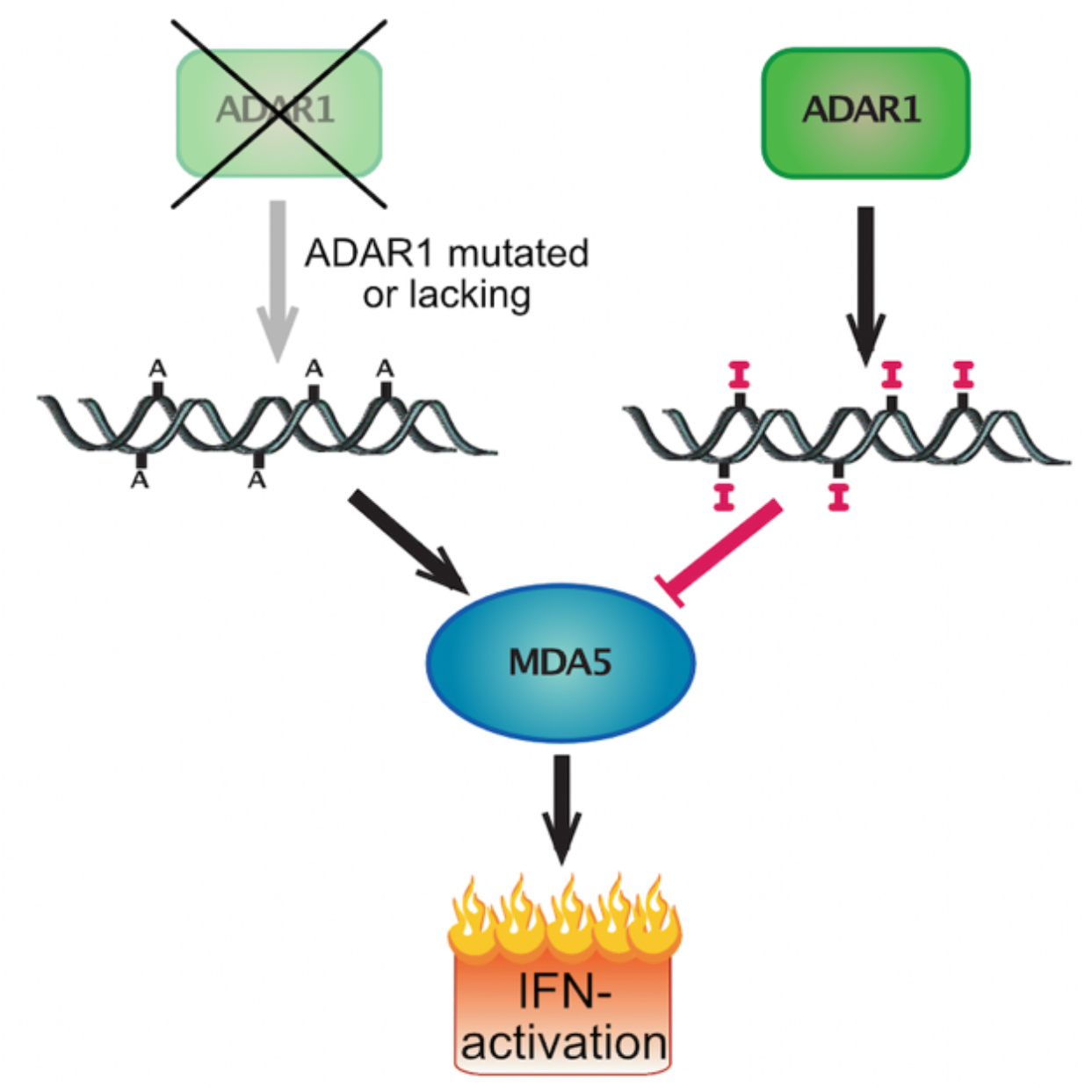
Sensing of dsRNA by MDA5: ADAR1 converts adenosines to inosines in dsRNA and thus labels RNAs as ’self’. In the absence of A-to-I editing, dsRNA triggers an innate immune response via activation of the cytoplasmic dsRNA sensor, MDA5.

**Figure 1 – figure supplement 3.**
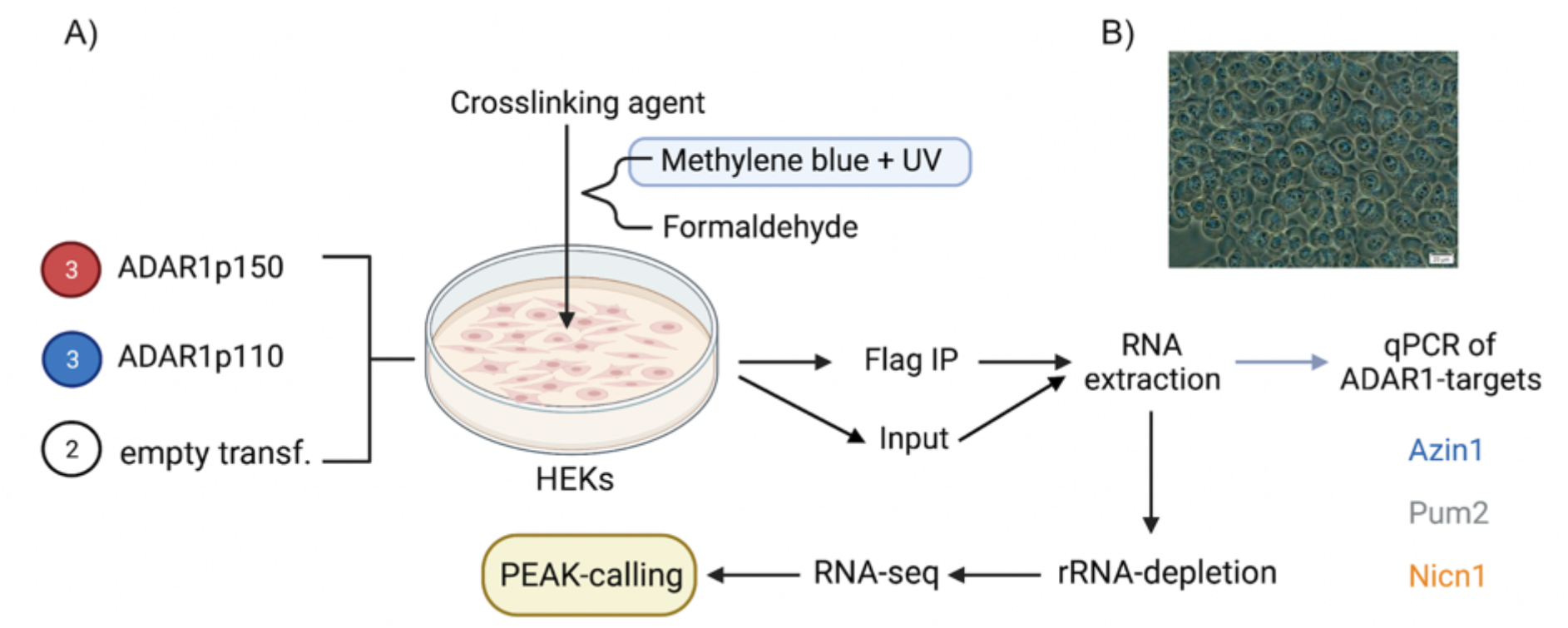
RIP-seq experimental workflow: A) HEK 293 cells were transfected with ADAR1p150-, ADAR1p110 – plasmid DNA or mock-empty transfection. The next day cells were subjected to selected cross-linking and harvested. The input sample was collected, and the remaining cell suspension was submitted to flag-IP. RNA was extracted from input- and IP- samples, and enrichment of selected editing substrates in IP fractions was examined using qPCR. Ribosomal RNA was depleted and RNA was converted to cDNA and subjected to high-throughput sequencing. The obtained reads were analysed via peak-calling. ADAR1p150 and ADAR1p110 RIP-seq were performed in triplicates and empty-mock transfection in duplicates. B) HEK 293 cells after exposure to methylene blue + UV cross-linking.

**Figure 1 – figure supplement 4.**
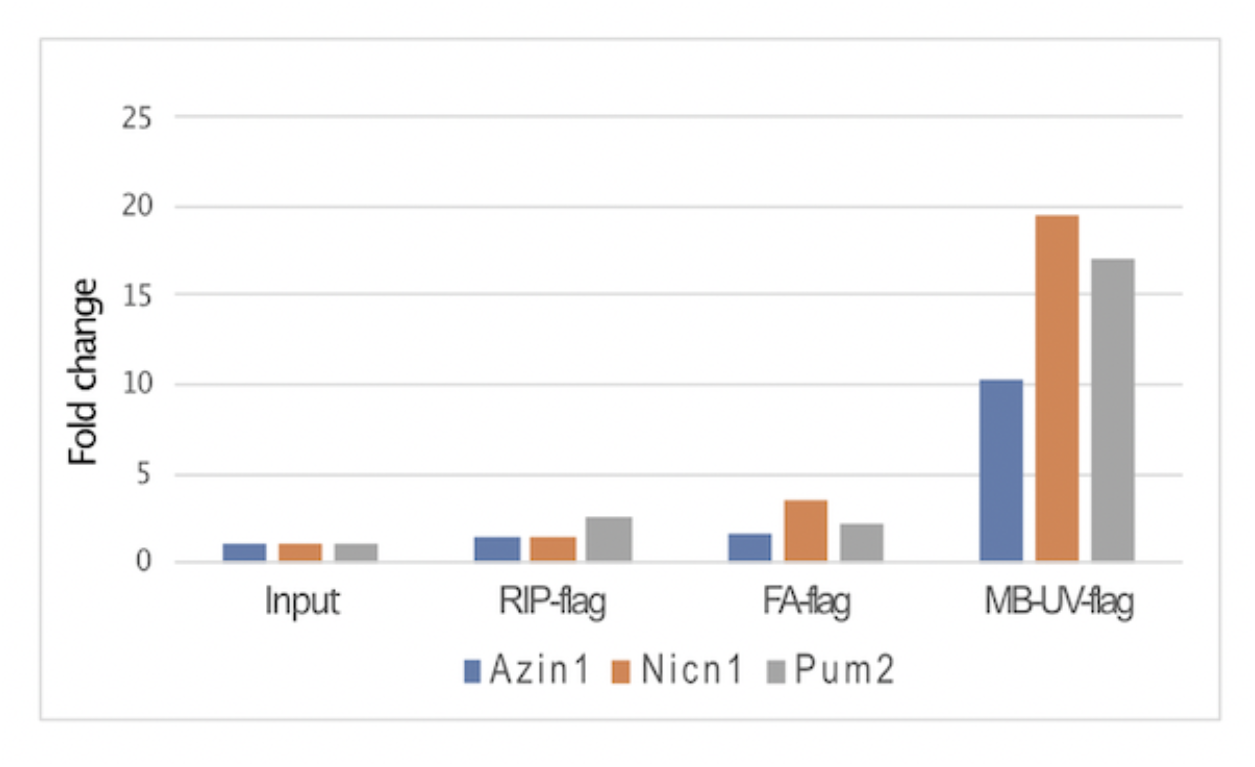
qPCR evaluation of ADAR1 targets in IP and corresponding input fraction upon selected cross-linking conditions. RIP-flag: no cross-linking applied; FA-flag: formaldehyde cross- linking; MB+UV-flag: a combination of methylene blue and ultra-violet light 254 nm. The experiment was conducted with the ADAR1p150 isoform. Fold change enrichment is normalised to the corresponding input.

**Figure 1 – figure supplement 5.**
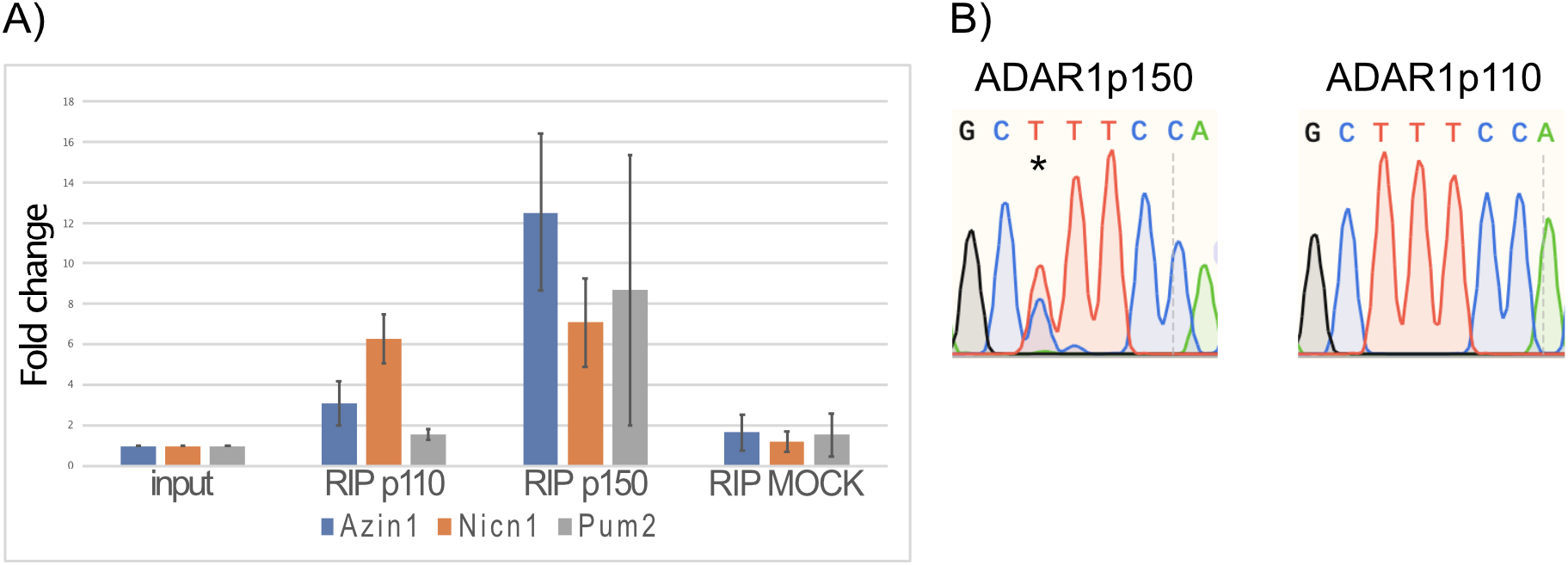
Verification of substrate enrichment and editing upon ADAR1 isoform IP: A) qPCR evaluation of enrichment of ADAR1-targets in IP-fractions. The enrichment was normalised to the relevant input sample. RIP p110 and RIP p150 were conducted in triplicates and RIP MOCK in duplicate. B) Sanger sequencing traces of a selected editing site. Azin1 (chr8:103841636) is primarily edited by ADAR1p150. The reverse strand is sequenced. Consequently, an A to I event is seen as a T to C conversion in the chromatogram-indicated with an asterisk (*)

**Figure 1 – figure supplement 6.**
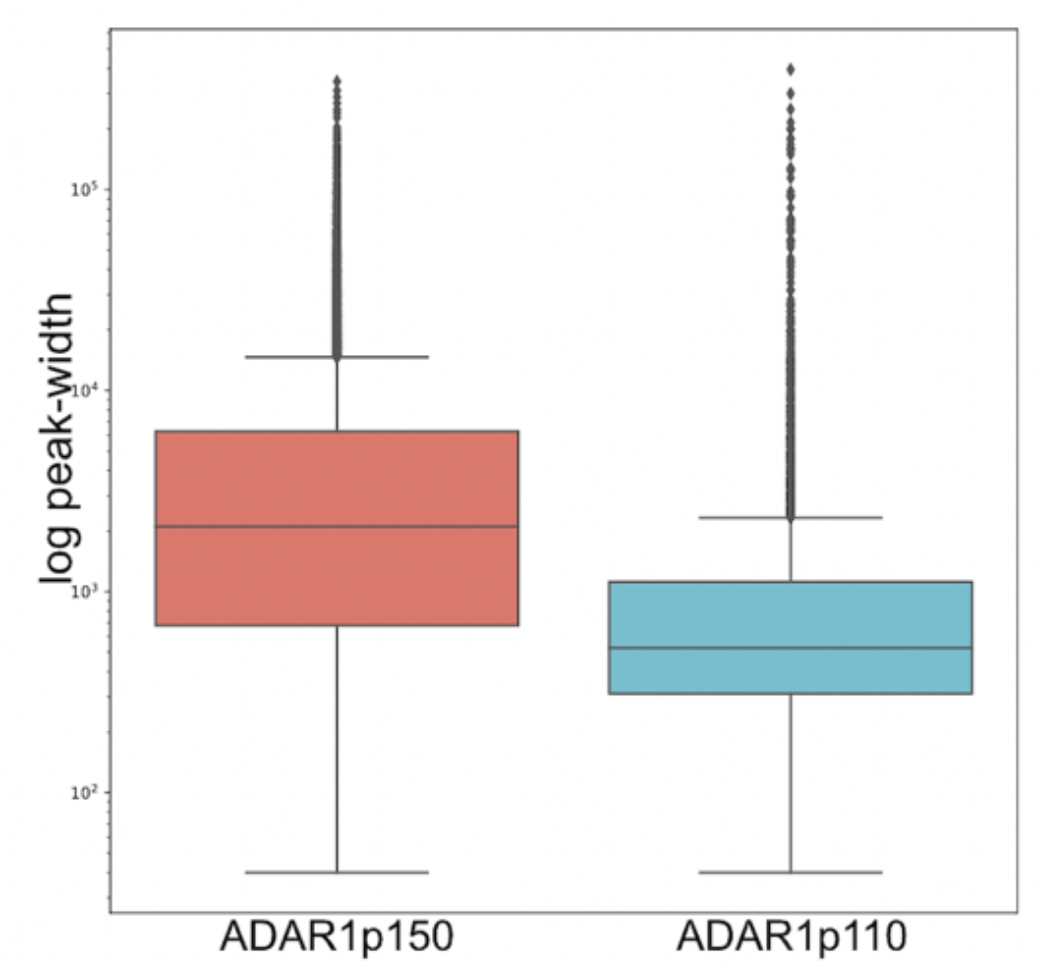
Peak-width distribution in RIP-seq experiments. Peak-size is based on genomic coordinates, and thus ADAR1p150-peaks are seemingly longer (median: 2105 nt) than ADAR1p110-peaks (median: 524 nt).

**Figure 1 – figure supplement 7.**
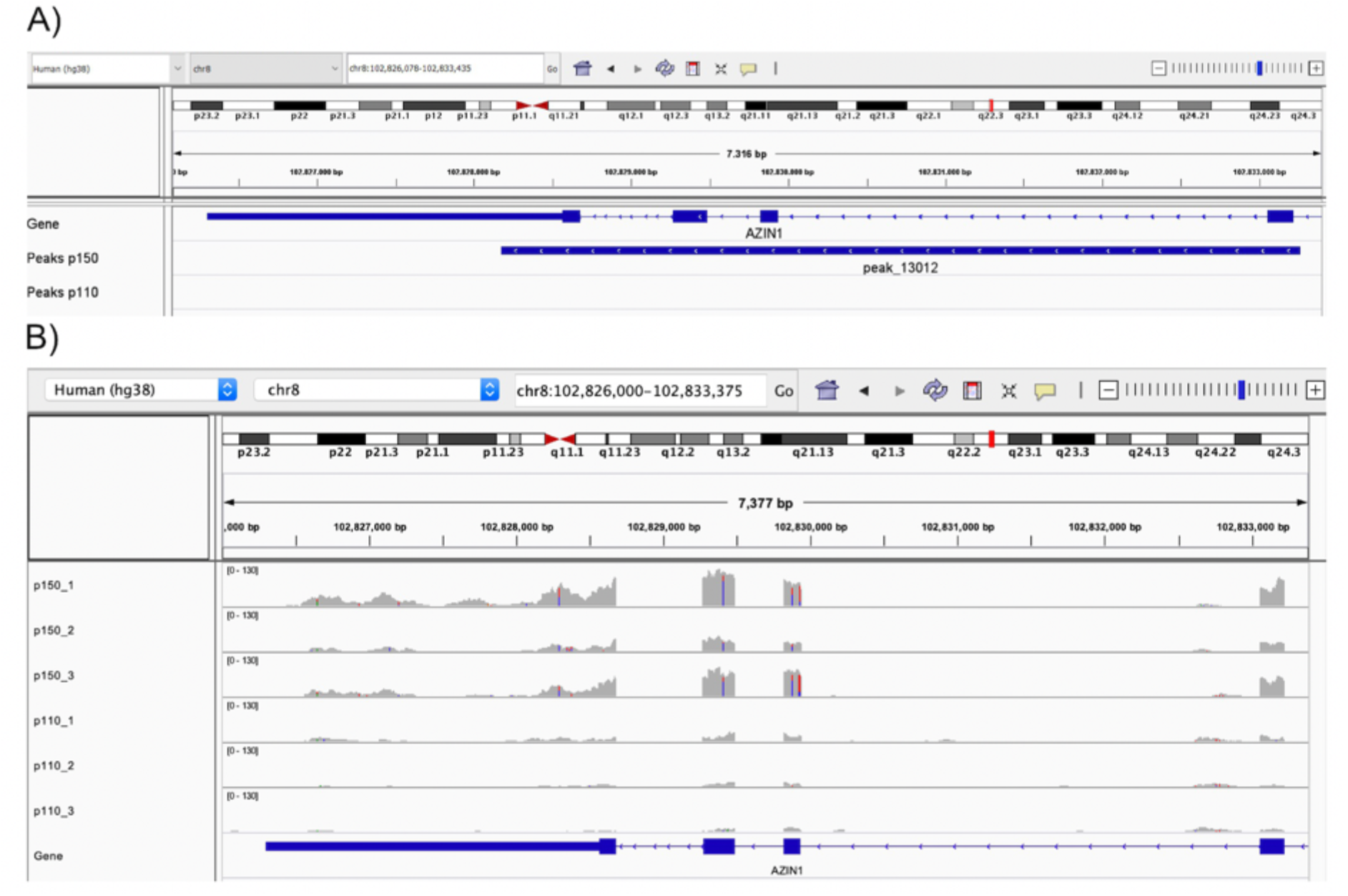
Peaks can span several exons. A) The peak 13012 spans several exons of Azin1. B) Read-coverage of IP samples in the peak-spanning region. Peak 13012 is clearly specific for ADAR1p150. The peak is physically formed almost exclusively from exons; therefore, the actual size of the peak is dramatically smaller than the size calculated based on the genomic coordinates.

**Figure 2 – figure supplement 1.**
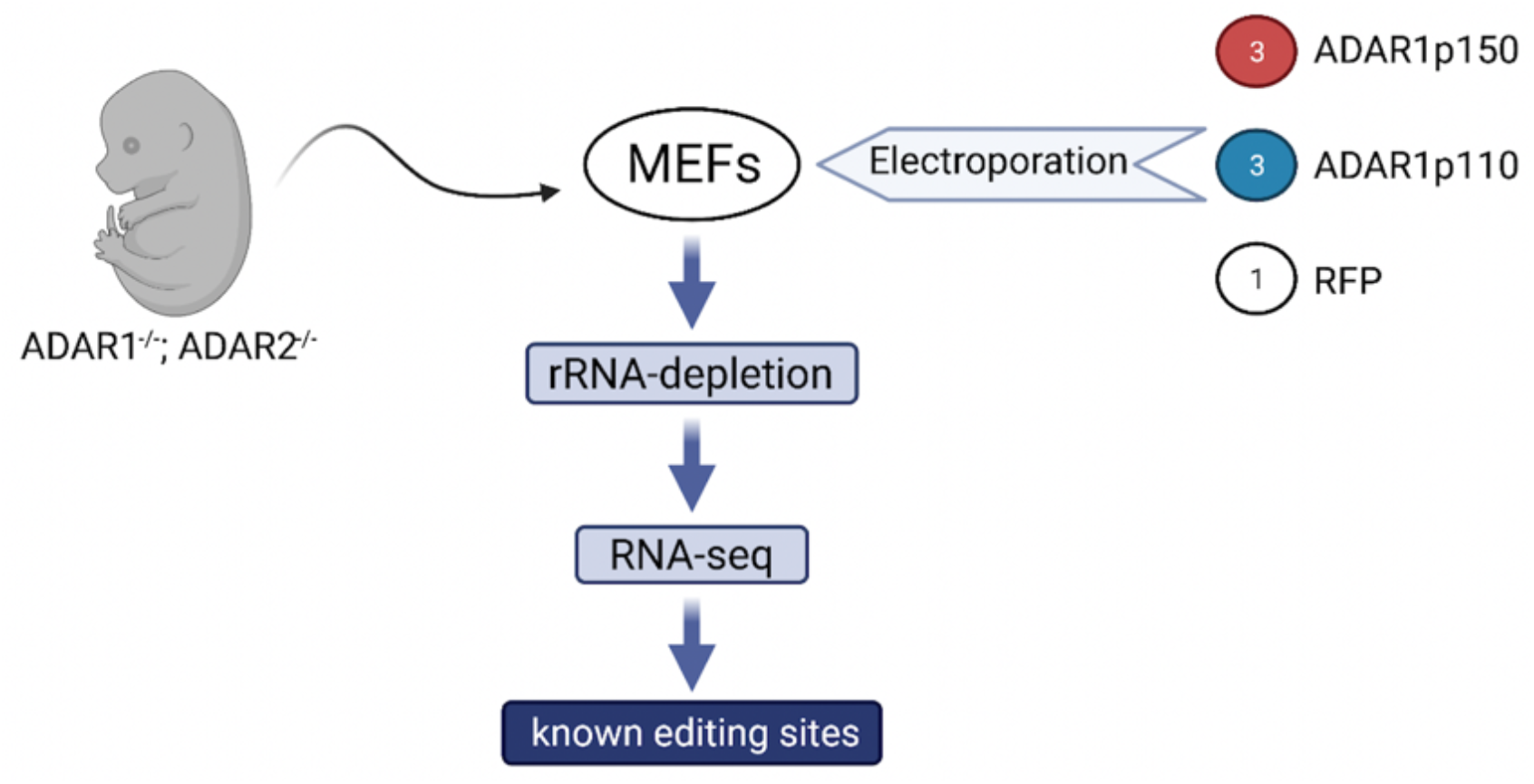
Experimental workflow of restoration of ADAR1 expression in editing deficient cells. Mouse embryonic fibroblasts (MEFs) generated from ADAR1^-/-^, ADAR2^-/-^ knockout mouse embryos, were electroporated with ADAR1p150, ADAR1p110 or RFP mammalian expression vector. 24 hours later cells were harvested and RNA was extracted. rRNA-depleted RNA-seq libraries were prepared and subjected to NGS. Editing was examined at know-editing sites.

**Figure 2 – figure supplement 2.**
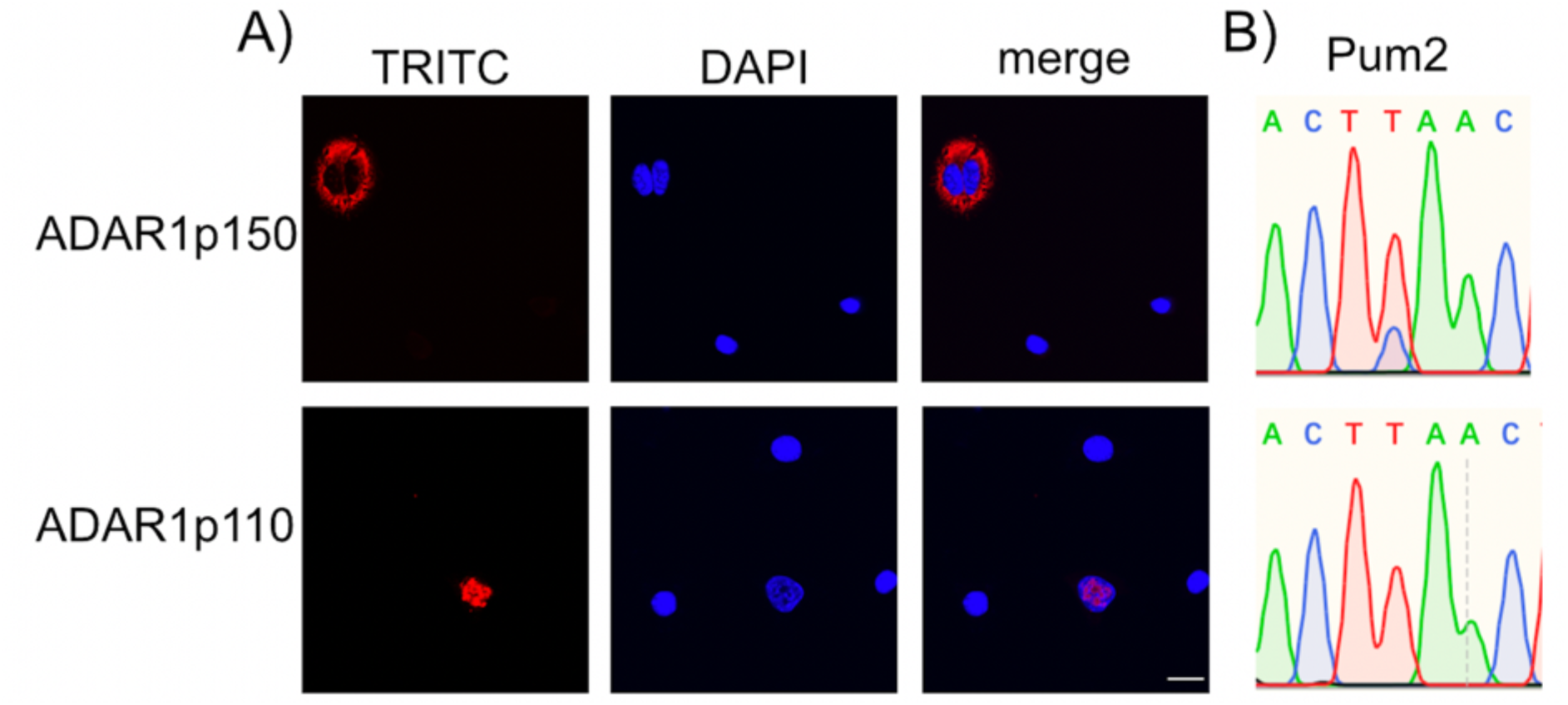
Over-expressed ADAR1-isoforms showed typical cellular localisation and distinct editing patterns. A) ADAR1p150 is mainly localised to the cytoplasm, whereas ADAR1p110 is primarily nuclear. TRITC channel shows transfected constructs in confocal sections, and nuclear DNA is stained with DAPI (Scale bar: 20 µm). B) Editing site in the 3’UTR of Pum2 (chr12: 8750269) is edited by ADAR1p150 but not by ADAR1p110.

**Figure 2 – figure supplement 3.**
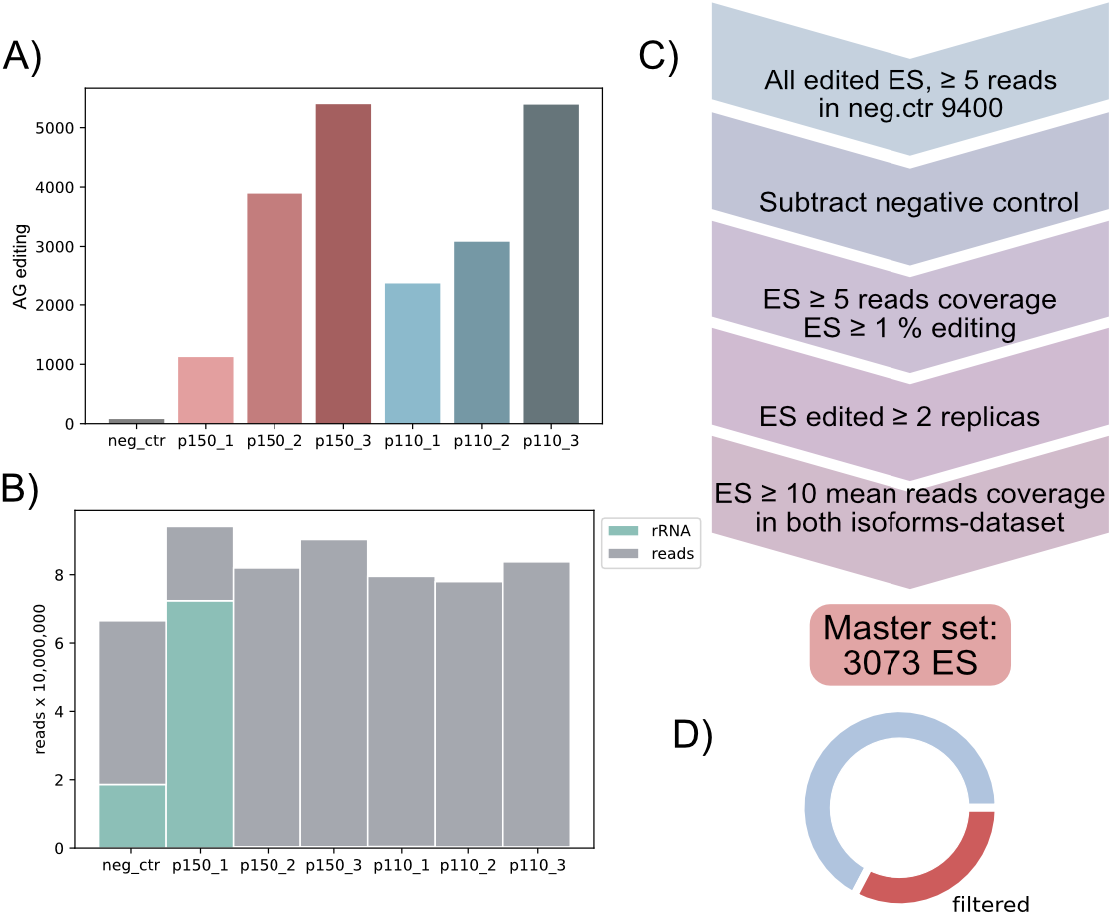
Restoring ADAR1 expression in editing-deficient cells produces an authentic set of editing sites. A) Editing in known-editing sites detected in each sample. Only few A to G transitions were detected in the negative control: 78 sites. Editing rises dramatically upon transfection with any ADAR1-isoform, ranging from 1121 to 5395 detected sites. Sites covered by ≥ 5 reads in at least two replicas showing ≥ 1% editing. B) rRNA reads in each sample. The first replica of ADAR1p150 shows a remarkably high rRNA fraction (77 %). While the negative control also has a prominent rRNA fraction the remaining samples exhibit a minimal rRNA portion (less than 1 %). C) Filtering strategy for the master set generation. All detected editing sites sufficiently covered (≥ 5 reads) in the negative control were combined, resulting in 9.400 sites. Next, sites found edited in the negative control were subtracted. Sites covered by at least 5 reads and showing an editing ratio of at least 1% in 2 out of 3 replicas were collected. Only sites with sufficient coverage in datasets of both isoforms (≥ 10 reads in average) formed the final ’master’ dataset containing 3.073 editing sites. D) Donut plot showing a portion of editing sites passing the filtering criteria.

**Figure 2 – figure supplement 4.**
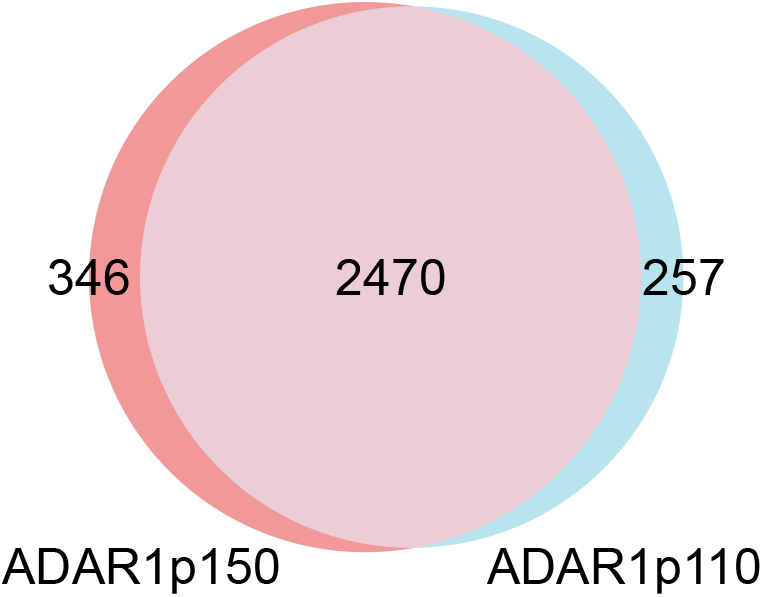
ADAR1 isoforms have a large overlap of editing sites: The majority of sites can be edited by both isoforms (also considering a minor editing rate below 1%). Still, 346 and 257 are exclusive for ADAR1p150 or ADAR1p110, respectively.

**Figure 3 – figure supplement 1.**
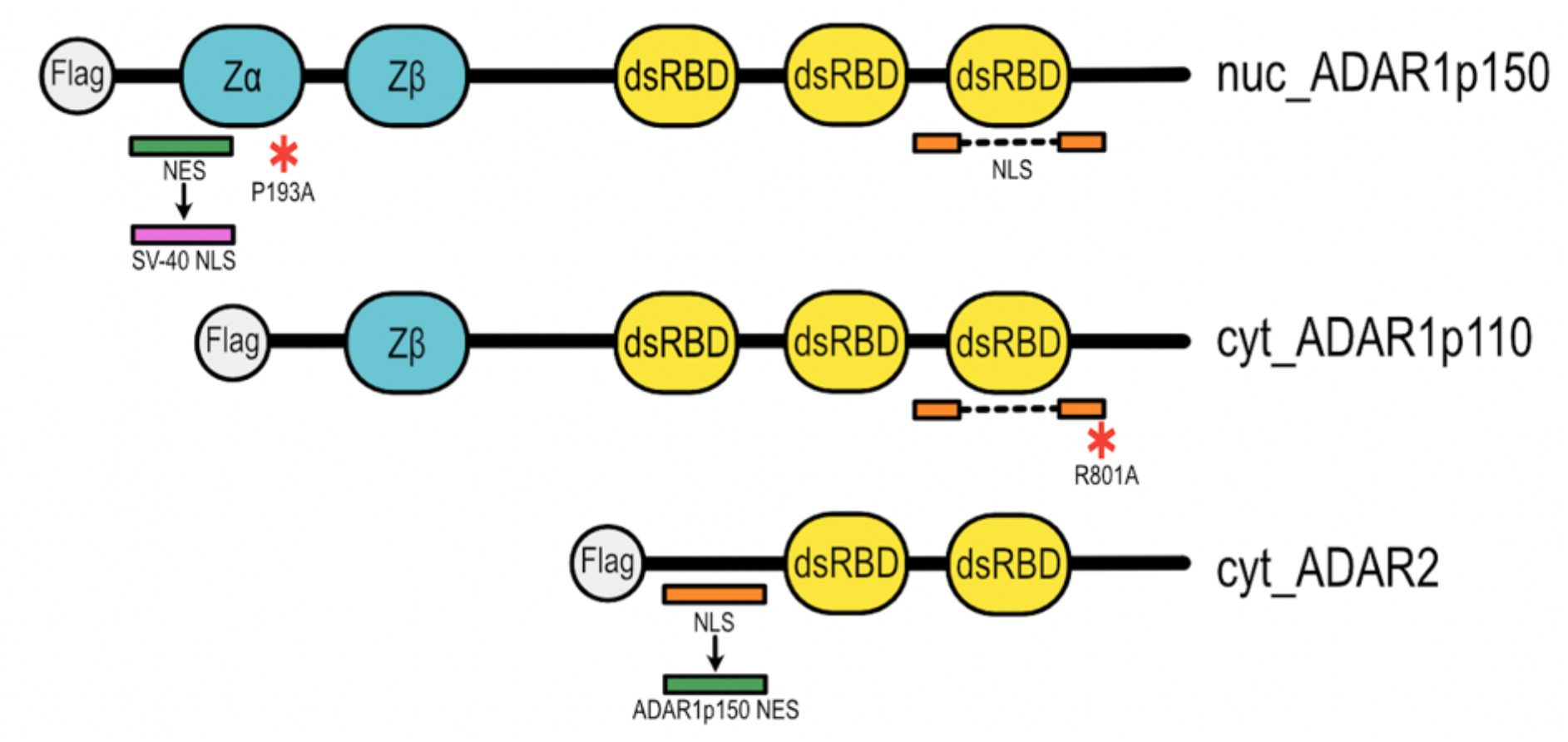
Scheme for the generation of mislocalized and ZBD*α* mutant ADARs. To construct nuclear ADAR1p150 the entire NES (CLSSHFQELSIYQDQEQRILKFLEEL, aa 125-150) was deleted and replaced with a SV-40 NLS (PKKKRKVEDP). The ZBD*α* mutant was generated by exchanging P193 to A193 to mimic the frequent mutation identified in AGS patients. Cytoplasmic ADAR1p110 was generated by introducing a point mutation: R801A that abolishes NLS activity. To prepare cytoplasmic ADAR2, the N-terminus was deleted (1-72) and replaced with the minimal NES of ADAR1p150 (CLSSHFQELSIY, 125-136)

**Figure 3 – figure supplement 2.**
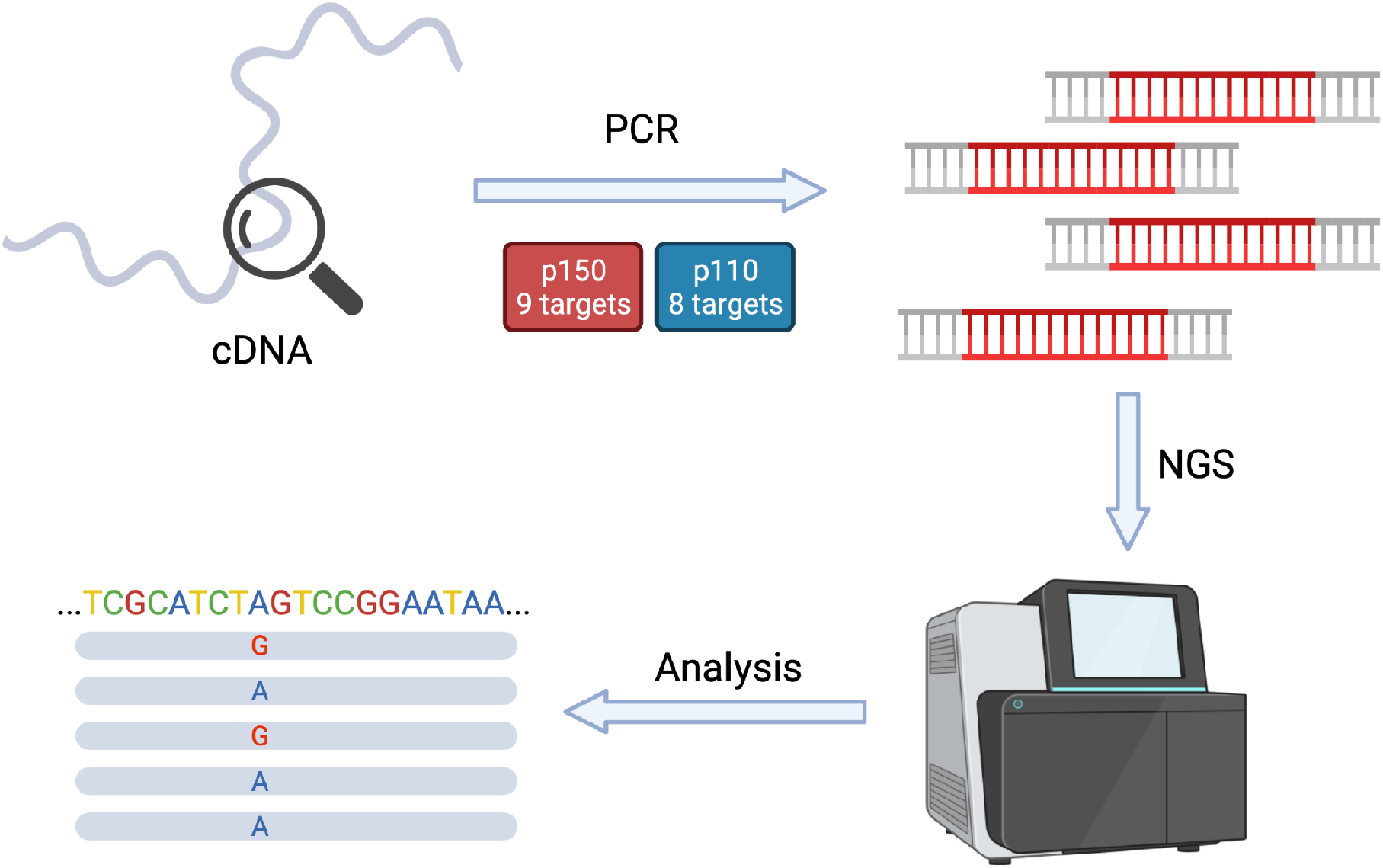
Workflow of amplicon-seq experiment: Total RNA was transcribed into cDNA. Amplicons were prepared via target-specific PCR with PCR bearing partial Illumina adaptor sequence. Next, amplicons were further amplified and barcoded by a second PCR reaction with primers containing complementary adaptor sequence and unique index-barcodes. Amplicons were sequenced, and the obtained reads were examined for editing events at known-editing sites.

**Figure 3 – figure supplement 3.**
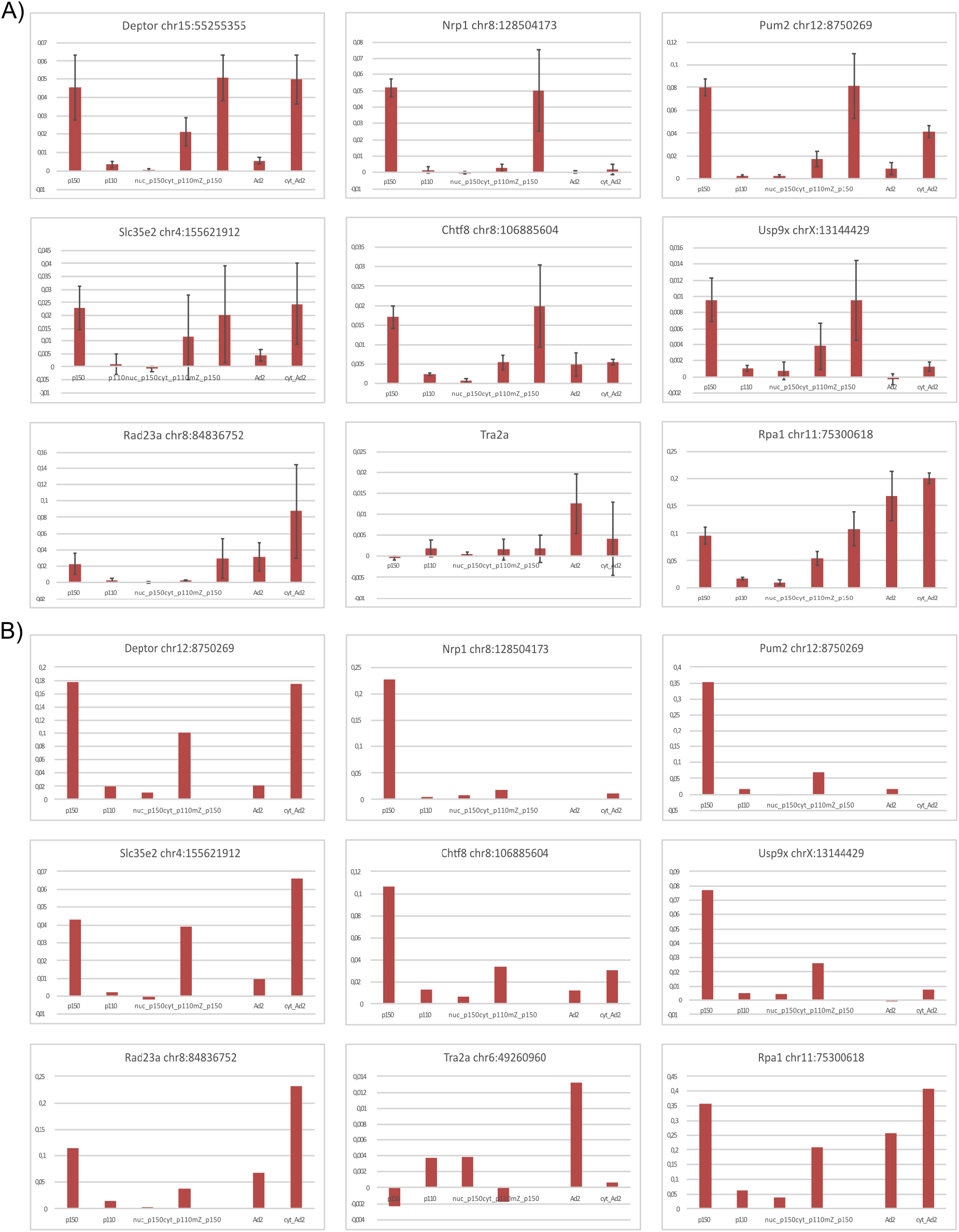
Editing detected by amplicon-seq in ADAR1p150-targets. The editing pattern remains similar upon transfection of 5µg and 10 µg of plasmid DNA A) Editing in the cells transfected with 5 µg of DNA; the experiment was conducted in triplicates; B) Editing in the cells transfected with 10 µg of DNA (this experiment does not contain mZ – ADAR1p150 P193A).

**Figure 3 – figure supplement 4.**
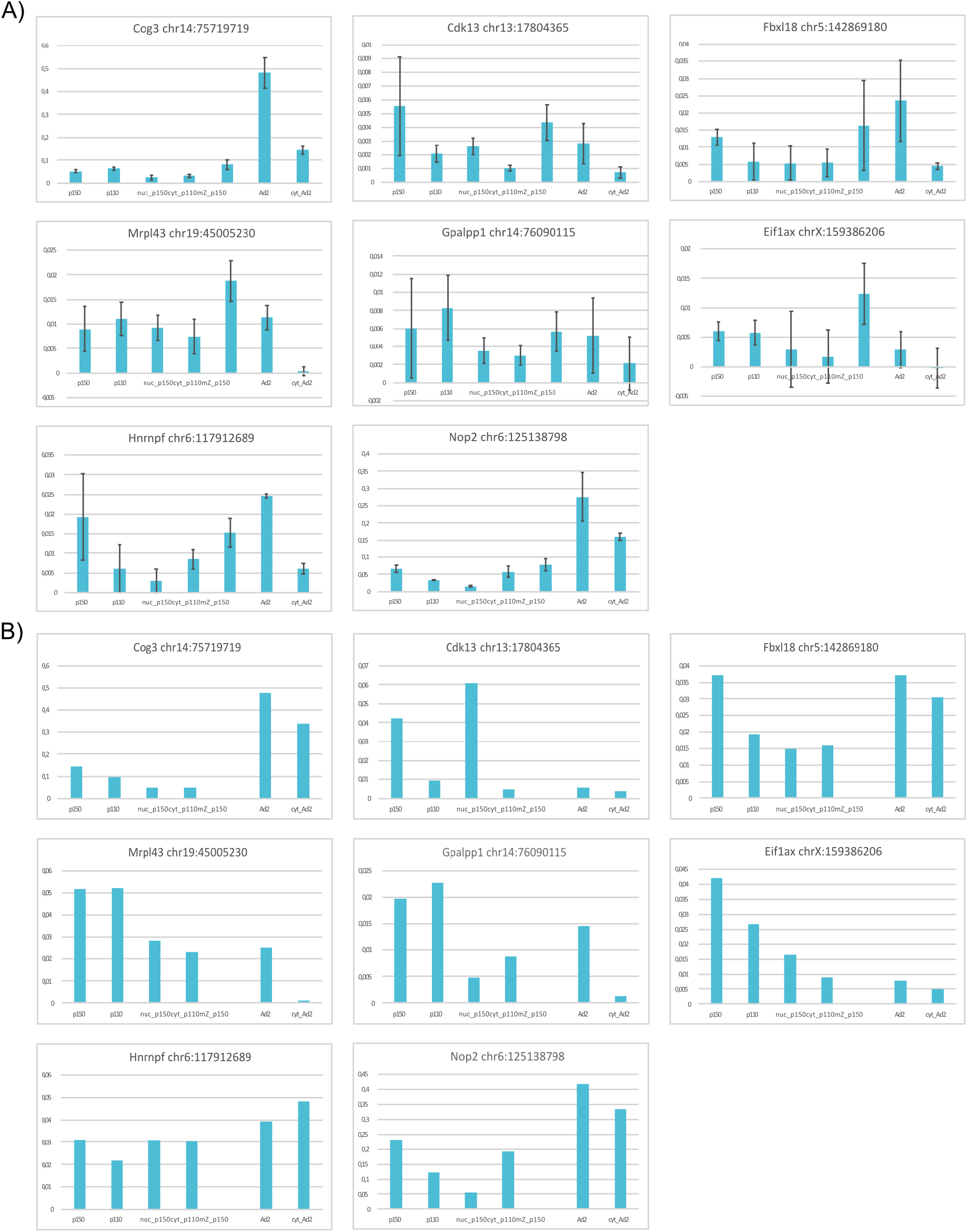
Editing detected by amplicon-seq in ADAR1p110-targets. The editing pattern remains similar upon transfection of 5µg and 10 µg of plasmid DNA. A) Editing in the cells transfected with 5 µg of DNA; the experiment was conducted in triplicates; B) Editing in the cells transfected with 10 µg of DNA (this experiment does not contain mZ – ADAR1p150 P193A).

**Figure 3 – figure supplement 5.**
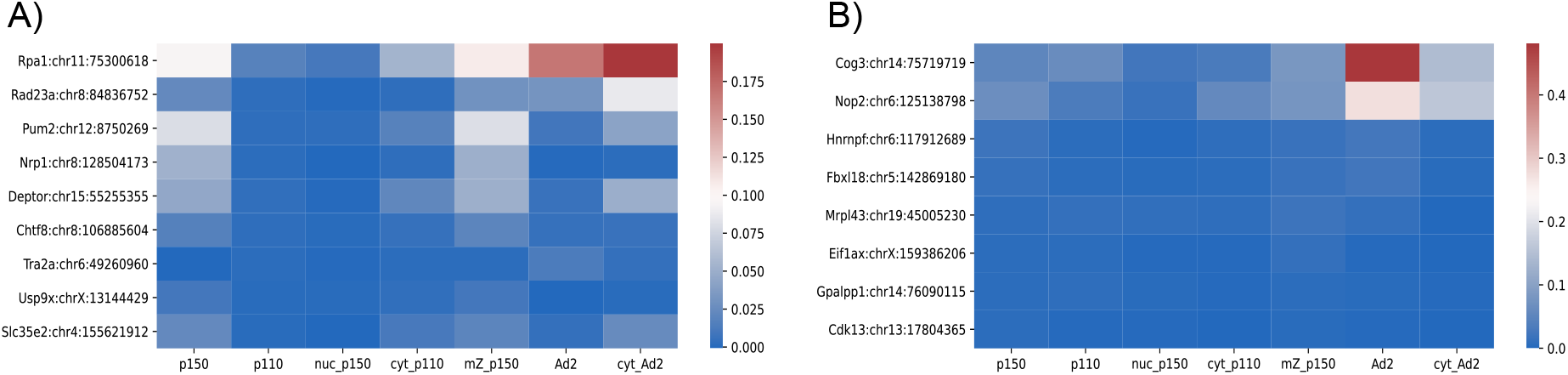
Heat maps for absolute editing ratios. A) Sites preferentially edited by ADAR1p150; B) Sites preferentially edited by ADAR1p110

**Figure 3 – figure supplement 6.**
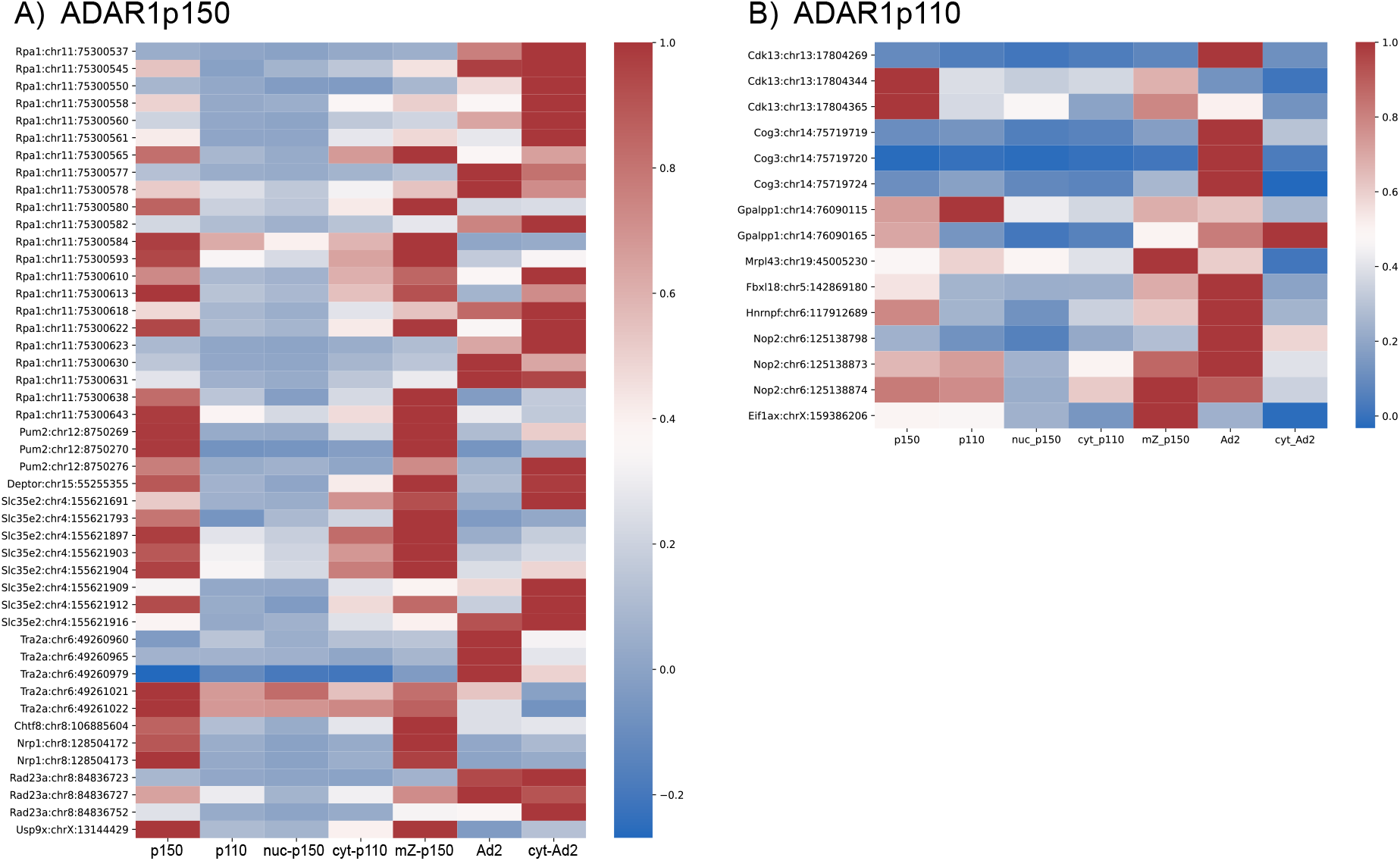
Heat maps of normalized editing ratio of all editing sites identified by amplicon-seq that are edited ≥ 0,5%. Apart from selected sites, amplicon-seq covers several closely located editing sites. Surprisingly, some very close sites show dramatically distinct editing patterns. A) Editing targets selected as preferentially edited by ADAR1p150 based on the ADAR1-editome obtained in MEFs; B) Editing targets selected as preferentially edited by ADAR1p110 based on the ADAR1-editome in MEFs.

**Figure 3 – figure supplement 7.**
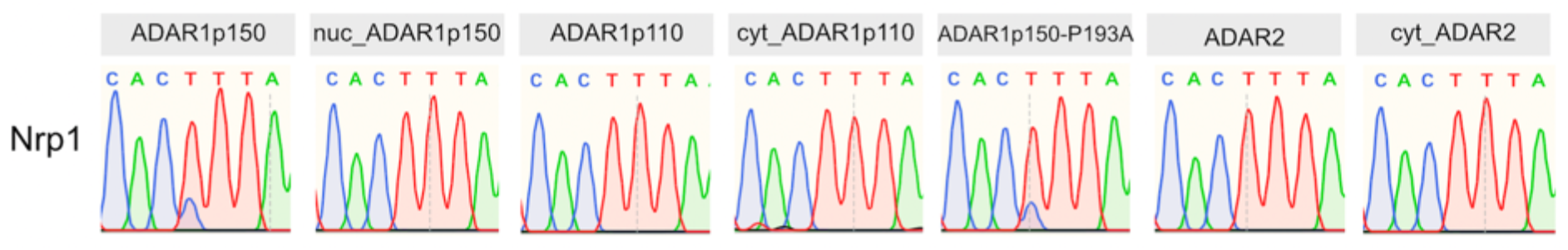
Sanger sequencing validates the Nrp1 editing pattern for wild-type and mutant ADAR isoforms as identified by amplicon-seq.

**Figure 4 – figure supplement 1.**
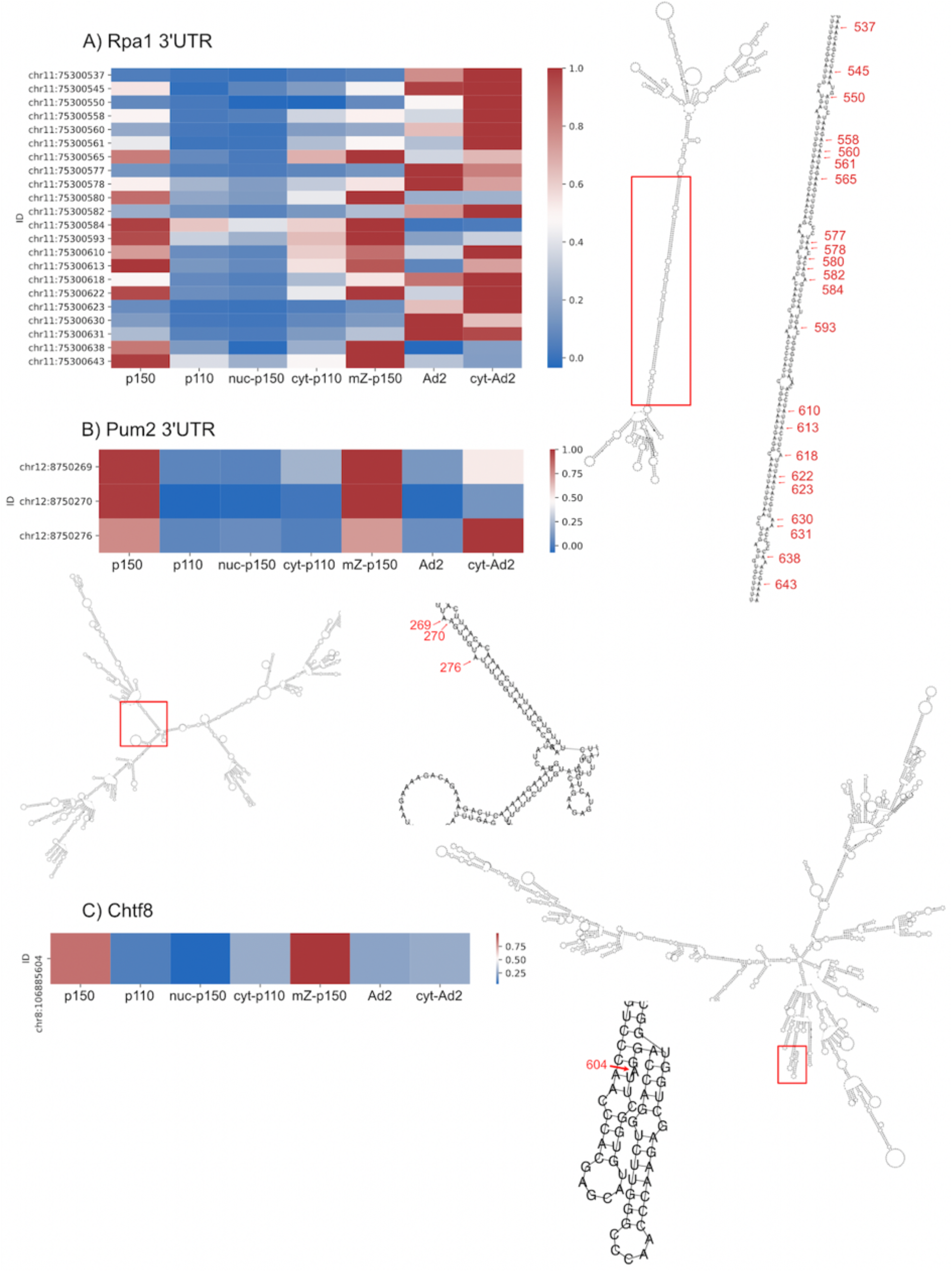
Heat maps of all editing sites detected by amplicon-seq in selected ADAR1p150 targets and folding prediction of substrates. Structure predictions were generated using Vienna RNA Package. Magnifications of edited regions display detected editing sites in the structure (labelled with the last 3 digits of the genomic coordinate).

**Figure 6 – figure supplement 1.**
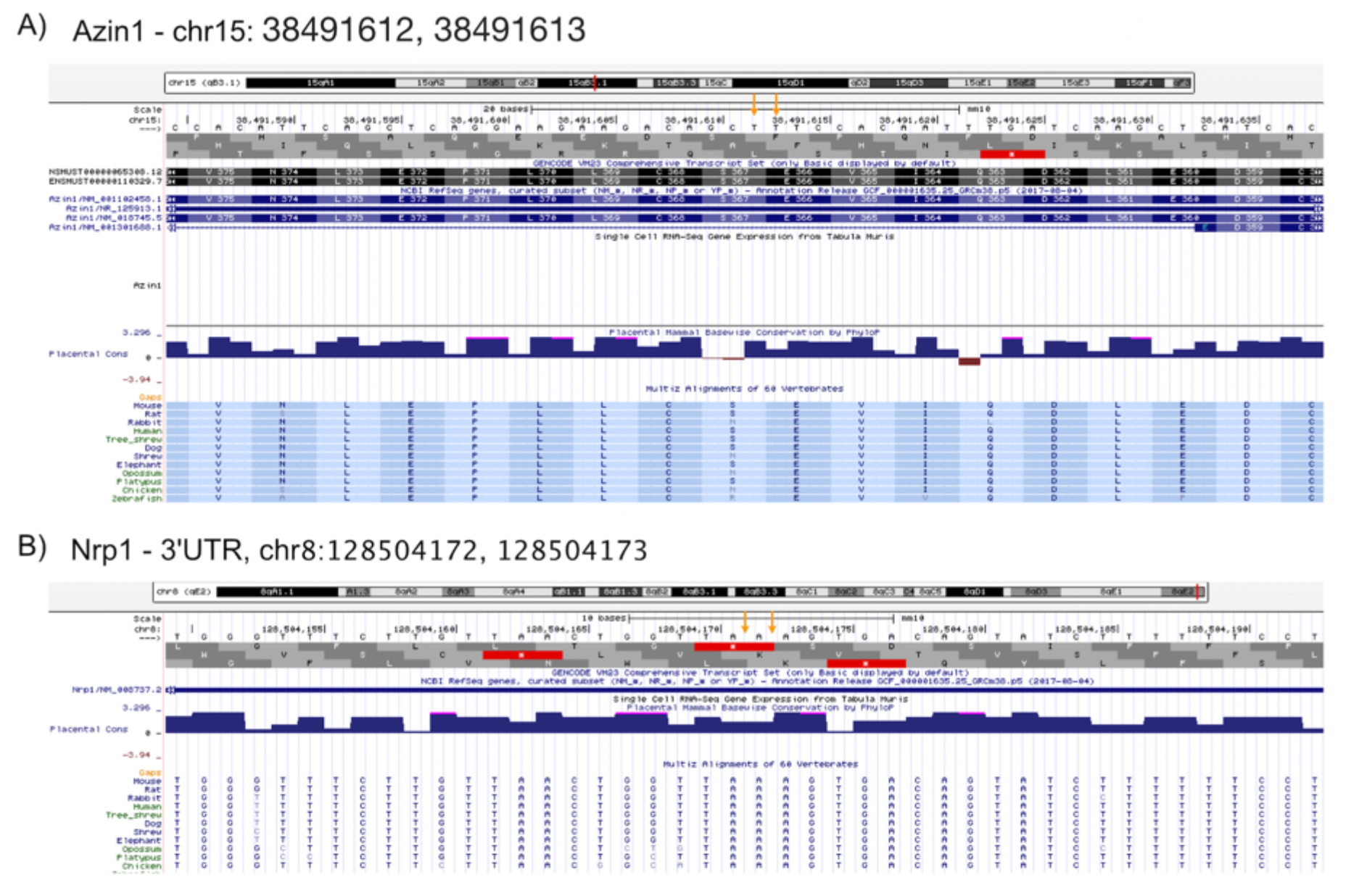
**Alignments generated with genome browser (http://genome.ucsc.edu) for selected editing sites.** A) Azin1 - chr15: 38491612, 38491613; B) Nrp1 - 3’UTR, chr8:128504172, 128504173.

**Figure 6 – figure supplement 2.**
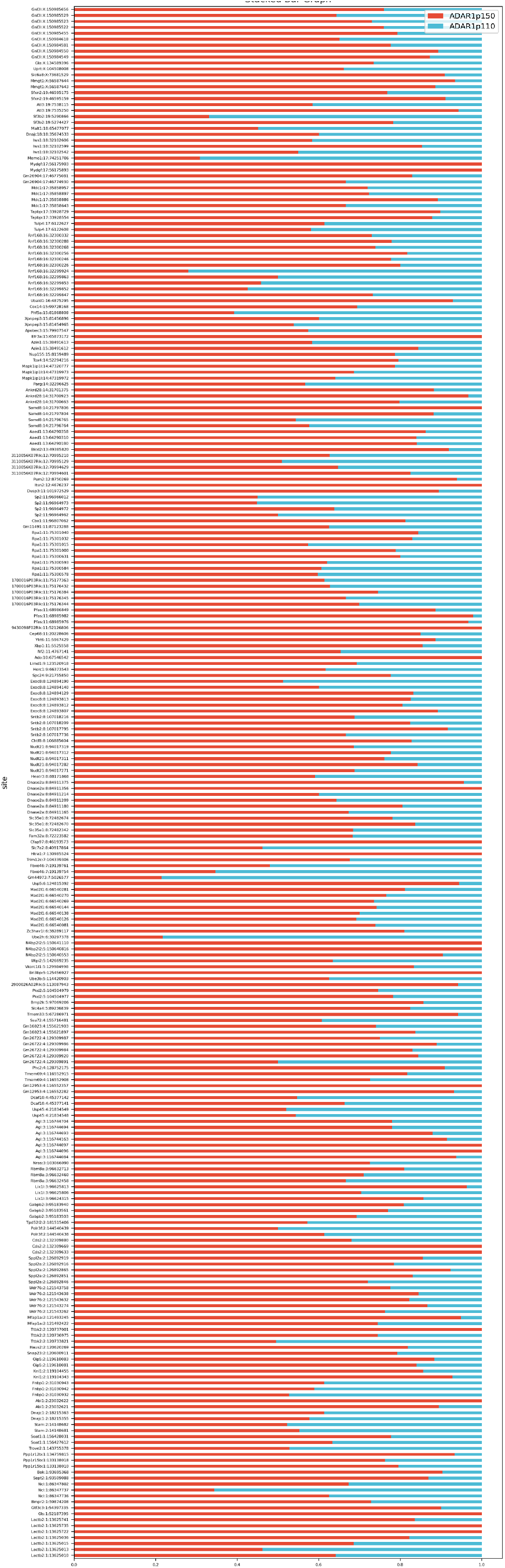
ADAR1p150 and ADAR1p110-mediated editing ratios detected in MEFs for sites found edited by ADAR1p150 in the thymus (Kim et al., 2021)

**Figure 6 – figure supplement 3.**
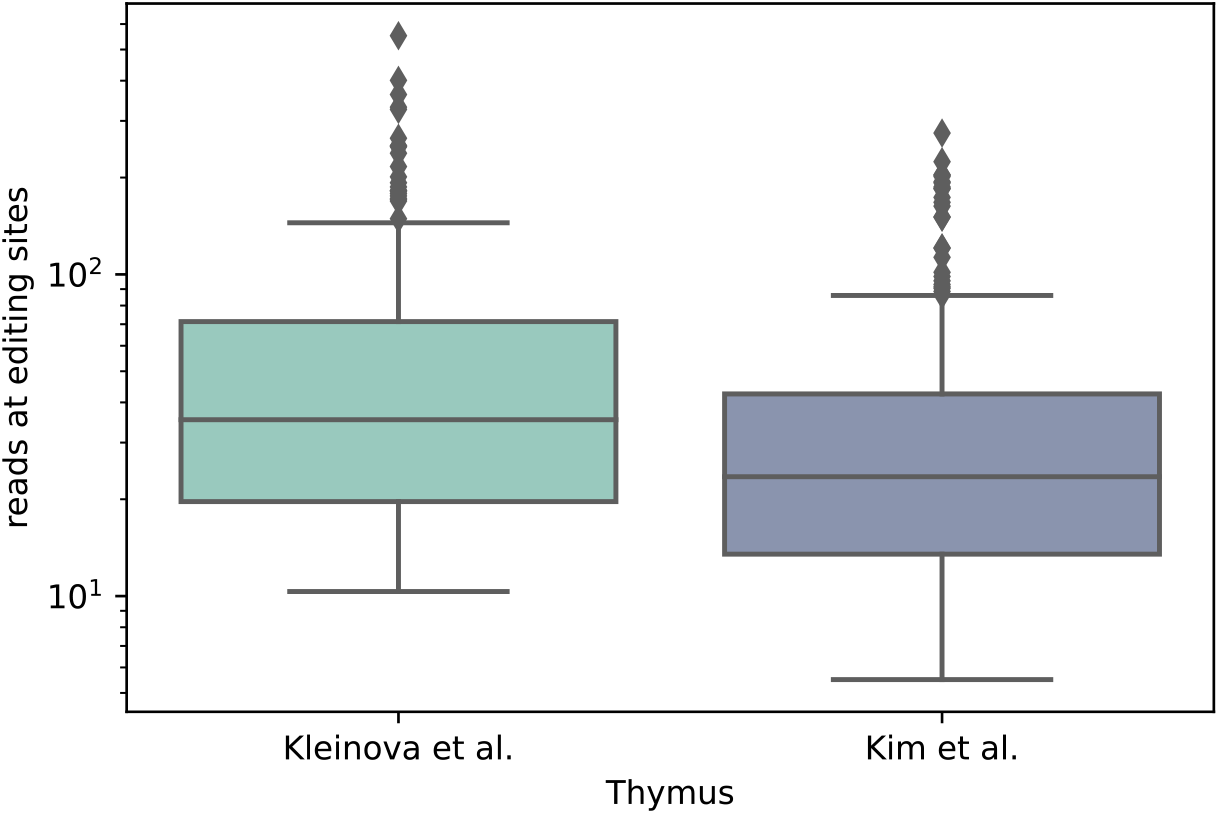
Read-coverage comparison of sites overlapping the MEF-editome analysis (this study- Kleinova et al.) and ADAR1p150-specific sites identified in the thymus by Kim et al., (Kim et al., 2021). The coverage of overlapping sites is almost 2-fold lower in the thymus sample.

## References

Afgan, E., Baker, D., Batut, B., van den Beek, M., Bouvier, D., Čech, M., Chilton, J., Clements, D., Coraor, N., Grüning, B.A., et al. (2018). The Galaxy platform for accessible, reproducible and collaborative biomedical analyses: 2018 update. Nucleic Acids Research 46, W537–W544. 10.1093/nar/gky379.

Ahmad, S., Mu, X., Yang, F., Greenwald, E., Park, J.W., Jacob, E., Zhang, C.Z., and Hur, S. (2018). Breaching Self-Tolerance to Alu Duplex RNA Underlies MDA5-Mediated Inflammation. Cell 172, 797–810 e713. 10.1016/j.cell.2017.12.016.

Barraud, P., Banerjee, S., Mohamed, W.I., Jantsch, M.F., and Allain, F.H. (2014). A bimodular nuclear localization signal assembled via an extended double-stranded RNA-binding domain acts as an RNA-sensing signal for transportin 1. Proc Natl Acad Sci U S A 111, E1852–1861. 10.1073/pnas.1323698111.

Bazak, L., Haviv, A., Barak, M., Jacob-Hirsch, J., Deng, P., Zhang, R., Isaacs, F.J., Rechavi, G., Li, J.B., Eisenberg, E., and Levanon, E.Y. (2014a). A-to-I RNA editing occurs at over a hundred million genomic sites, located in a majority of human genes. Genome Res 24, 365–376. 10.1101/gr.164749.113.

Bazak, L., Levanon, E.Y., and Eisenberg, E. (2014b). Genome-wide analysis of Alu editability. Nucleic Acids Res 42, 6876–6884. 10.1093/nar/gku414.

Behm, M., Wahlstedt, H., Widmark, A., Eriksson, M., and Ohman, M. (2017). Accumulation of nuclear ADAR2 regulates adenosine-to-inosine RNA editing during neuronal development. J Cell Sci 130, 745–753. 10.1242/jcs.200055.

Bentley, D.L. (2014). Coupling mRNA processing with transcription in time and space. Nat Rev Genet 15, 163–175. 10.1038/nrg3662.

Brown, B.A., 2nd, Lowenhaupt, K., Wilbert, C.M., Hanlon, E.B., and Rich, A. (2000). The zalpha domain of the editing enzyme dsRNA adenosine deaminase binds left-handed Z-RNA as well as Z-DNA. Proc Natl Acad Sci U S A 97, 13532–13536. 10.1073/pnas.240464097 240464097 [pii].

Cantuti-Castelvetri, L., Ojha, R., Pedro, L.D., Djannatian, M., Franz, J., Kuivanen, S., van der Meer, F., Kallio, K., Kaya, T., Anastasina, M., et al. (2020). Neuropilin-1 facilitates SARS-CoV-2 cell entry and infectivity. Science 370, 856–860. 10.1126/science.abd2985.

Chen, S., Zhou, Y., Chen, Y., and Gu, J. (2018). fastp: an ultra-fast all-in-one FASTQ preprocessor. Bioinformatics 34, i884–i890. 10.1093/bioinformatics/bty560.

Crow, Y.J., Chase, D.S., Lowenstein Schmidt, J., Szynkiewicz, M., Forte, G.M., Gornall, H.L., Oojageer, A., Anderson, B., Pizzino, A., Helman, G., et al. (2015). Characterization of human disease phenotypes associated with mutations in TREX1, RNASEH2A, RNASEH2B, RNASEH2C, SAMHD1, ADAR, and IFIH1. Am J Med Genet A 167A, 296–312. 10.1002/ajmg.a.36887.

de Reuver, R., Dierick, E., Wiernicki, B., Staes, K., Seys, L., De Meester, E., Muyldermans, T., Botzki, A., Lambrecht, B.N., Van Nieuwerburgh, F., et al. (2021). ADAR1 interaction with Z- RNA promotes editing of endogenous double-stranded RNA and prevents MDA5-dependent immune activation. Cell reports 36, 109500. 10.1016/j.celrep.2021.109500.

Desterro, J.M., Keegan, L.P., Lafarga, M., Berciano, M.T., O’Connell, M., and Carmo-Fonseca, M. (2003). Dynamic association of RNA-editing enzymes with the nucleolus. J Cell Sci 116, 1805–1818. 10.1242/jcs.00371.

Dobin, A., Davis, C.A., Schlesinger, F., Drenkow, J., Zaleski, C., Jha, S., Batut, P., Chaisson, M., and Gingeras, T.R. (2012). STAR: ultrafast universal RNA-seq aligner. Bioinformatics 29, 15–21. 10.1093/bioinformatics/bts635.

Dobin, A., Davis, C.A., Schlesinger, F., Drenkow, J., Zaleski, C., Jha, S., Batut, P., Chaisson, M., and Gingeras, T.R. (2013). STAR: ultrafast universal RNA-seq aligner. Bioinformatics 29, 15–21. 10.1093/bioinformatics/bts635.

Eckmann, C.R., Neunteufl, A., Pfaffstetter, L., and Jantsch, M.F. (2001). The human but not the Xenopus RNA-editing enzyme ADAR1 has an atypical nuclear localization signal and displays the characteristics of a shuttling protein. Mol Biol Cell 12, 1911–1924.

Faulkner, G.J., Kimura, Y., Daub, C.O., Wani, S., Plessy, C., Irvine, K.M., Schroder, K., Cloonan, N., Steptoe, A.L., Lassmann, T., et al. (2009). The regulated retrotransposon transcriptome of mammalian cells. Nat Genet 41, 563–571. 10.1038/ng.368.

Fritz, J., Strehblow, A., Taschner, A., Schopoff, S., Pasierbek, P., and Jantsch, M.F. (2009). RNA-regulated interaction of transportin-1 and exportin-5 with the double-stranded RNA- binding domain regulates nucleocytoplasmic shuttling of ADAR1. Mol Cell Biol 29, 1487–1497. 10.1128/MCB.01519-08.

George, C.X., and Samuel, C.E. (1999). Human RNA-specific adenosine deaminase ADAR1 transcripts possess alternative exon 1 structures that initiate from different promoters, one constitutively active and the other interferon inducible. Proc Natl Acad Sci U S A 96, 4621–4626. 10.1073/pnas.96.8.4621.

Gruber, A.R., Lorenz, R., Bernhart, S.H., Neubock, R., and Hofacker, I.L. (2008). The Vienna RNA websuite. Nucleic Acids Res 36, W70–74. 10.1093/nar/gkn188.

Hartner, J.C., Schmittwolf, C., Kispert, A., Muller, A.M., Higuchi, M., and Seeburg, P.H. (2004). Liver disintegration in the mouse embryo caused by deficiency in the RNA-editing enzyme ADAR1. J Biol Chem 279, 4894–4902. 10.1074/jbc.M311347200.

Herbert, A. (2019). Z-DNA and Z-RNA in human disease. Commun Biol 2, 7. 10.1038/s42003-018-0237-x.

Herbert, A., and Rich, A. (1999). Left-handed Z-DNA: structure and function. Genetica 106, 37–47.

Higuchi, M., Maas, S., Single, F.N., Hartner, J., Rozov, A., Burnashev, N., Feldmeyer, D., Sprengel, R., and Seeburg, P.H. (2000). Point mutation in an AMPA receptor gene rescues lethality in mice deficient in the RNA-editing enzyme ADAR2. Nature 406, 78–81. 10.1038/35017558.

Hsieh, C.L., Liu, H., Huang, Y., Kang, L., Chen, H.W., Chen, Y.T., Wee, Y.R., Chen, S.J., and Tan, B.C. (2014). ADAR1 deaminase contributes to scheduled skeletal myogenesis progression via stage-specific functions. Cell Death Differ 21, 707–719. 10.1038/cdd.2013.197.

Hu, S., Hu, Z., Qin, J., Lin, C., and Jiang, X. (2021). In silico analysis identifies neuropilin-1 as a potential therapeutic target for SARS-Cov-2 infected lung cancer patients. Aging (Albany NY) 13, 15770–15784. 10.18632/aging.203159.

Hu, X., Chen, J., Shi, X., Feng, F., Lau, K.W., Chen, Y., Chen, Y., Jiang, L., Cui, F., Zhang, Y., et al. (2017). RNA editing of AZIN1 induces the malignant progression of non-small-cell lung cancers. Tumour Biol 39, 1010428317700001. 10.1177/1010428317700001.

Ishizuka, J.J., Manguso, R.T., Cheruiyot, C.K., Bi, K., Panda, A., Iracheta-Vellve, A., Miller, B.C., Du, P.P., Yates, K.B., Dubrot, J., et al. (2018). Loss of ADAR1 in tumours overcomes resistance to immune checkpoint blockade. Nature. 10.1038/s41586-018-0768-9.

Kim, J.I., Nakahama, T., Yamasaki, R., Costa Cruz, P.H., Vongpipatana, T., Inoue, M., Kanou, N., Xing, Y., Todo, H., Shibuya, T., et al. (2021). RNA editing at a limited number of sites is sufficient to prevent MDA5 activation in the mouse brain. PLoS Genet 17, e1009516. 10.1371/journal.pgen.1009516.

Koeris, M., Funke, L., Shrestha, J., Rich, A., and Maas, S. (2005). Modulation of ADAR1 editing activity by Z-RNA in vitro. Nucleic Acids Res 33, 5362–5370. 10.1093/nar/gki849.

Li, H., Handsaker, B., Wysoker, A., Fennell, T., Ruan, J., Homer, N., Marth, G., Abecasis, G., Durbin, R., and Genome Project Data Processing, S. (2009). The Sequence Alignment/Map format and SAMtools. Bioinformatics 25, 2078–2079. 10.1093/bioinformatics/btp352.

Licht, K., Kapoor, U., Amman, F., Picardi, E., Martin, D., Bajad, P., and Jantsch, M.F. (2019). A high resolution A-to-I editing map in the mouse identifies editing events controlled by pre- mRNA splicing. Genome Res 29, 1453–1463. 10.1101/gr.242636.118.

Licht, K., Kapoor, U., Mayrhofer, E., and Jantsch, M.F. (2016). Adenosine to Inosine editing frequency controlled by splicing efficiency. Nucleic Acids Res 44, 6398–6408. 10.1093/nar/gkw325.

Liddicoat, B.J., Piskol, R., Chalk, A.M., Ramaswami, G., Higuchi, M., Hartner, J.C., Li, J.B., Seeburg, P.H., and Walkley, C.R. (2015). RNA editing by ADAR1 prevents MDA5 sensing of endogenous dsRNA as nonself. Science 349, 1115–1120. 10.1126/science.aac7049.

Lin, C., and Miles, W.O. (2019). Beyond CLIP: advances and opportunities to measure RBP- RNA and RNA-RNA interactions. Nucleic Acids Res 47, 5490–5501. 10.1093/nar/gkz295.

Liu, Z.R., Wilkie, A.M., Clemens, M.J., and Smith, C.W. (1996). Detection of double-stranded RNA-protein interactions by methylene blue-mediated photo-crosslinking. RNA 2, 611–621.

Livingston, J.H., Lin, J.P., Dale, R.C., Gill, D., Brogan, P., Munnich, A., Kurian, M.A., Gonzalez- Martinez, V., De Goede, C.G., Falconer, A., et al. (2014). A type I interferon signature identifies bilateral striatal necrosis due to mutations in ADAR1. J Med Genet 51, 76–82. 10.1136/jmedgenet-2013-102038.

Lo Giudice, C., Tangaro, M.A., Pesole, G., and Picardi, E. (2020). Investigating RNA editing in deep transcriptome datasets with REDItools and REDIportal. Nat Protoc 15, 1098–1131. 10.1038/s41596-019-0279-7.

Lorenz, R., Bernhart, S.H., Honer Zu Siederdissen, C., Tafer, H., Flamm, C., Stadler, P.F., and Hofacker, I.L. (2011). ViennaRNA Package 2.0. Algorithms Mol Biol 6, 26. 10.1186/1748-7188-6-26.

Mannion, N.M., Greenwood, S.M., Young, R., Cox, S., Brindle, J., Read, D., Nellaker, C., Vesely, C., Ponting, C.P., McLaughlin, P.J., et al. (2014). The RNA-editing enzyme ADAR1 controls innate immune responses to RNA. Cell reports 9, 1482–1494. 10.1016/j.celrep.2014.10.041.

Martin, M. (2011). Cutadapt removes adapter sequences from high-throughput sequencing reads. 2011 *17*, 3. 10.14806/ej.17.1.200.

Maurano, M., Snyder, J.M., Connelly, C., Henao-Mejia, J., Sidrauski, C., and Stetson, D.B. (2021a). Protein kinase R and the integrated stress response drive immunopathology caused by mutations in the RNA deaminase ADAR1. Immunity 54, 1948–1960 e1945. 10.1016/j.immuni.2021.07.001.

Maurano, M., Snyder, J.M., Connelly, C., Henao-Mejia, J., Sidrauski, C., and Stetson, D.B. (2021b). Protein kinase R and the integrated stress response drive immunopathology caused by mutations in the RNA deaminase ADAR1. Immunity. 10.1016/j.immuni.2021.07.001.

Nakahama, T., Kato, Y., Shibuya, T., Inoue, M., Kim, J.I., Vongpipatana, T., Todo, H., Xing, Y., and Kawahara, Y. (2021). Mutations in the adenosine deaminase ADAR1 that prevent endogenous Z-RNA binding induce Aicardi-Goutieres-syndrome-like encephalopathy. Immunity 54, 1976–1988 e1977. 10.1016/j.immuni.2021.08.022.

Nishikura, K. (2010). Functions and regulation of RNA editing by ADAR deaminases. Annu Rev Biochem 79, 321–349. 10.1146/annurev-biochem-060208-105251.

Nishikura, K. (2016). A-to-I editing of coding and non-coding RNAs by ADARs. Nat Rev Mol Cell Biol 17, 83–96. 10.1038/nrm.2015.4.

Okugawa, Y., Toiyama, Y., Shigeyasu, K., Yamamoto, A., Shigemori, T., Yin, C., Ichikawa, T., Yasuda, H., Fujikawa, H., Yoshiyama, S., et al. (2018). Enhanced AZIN1 RNA editing and overexpression of its regulatory enzyme ADAR1 are important prognostic biomarkers in gastric cancer. J Transl Med 16, 366. 10.1186/s12967-018-1740-z.

Patterson, J.B., and Samuel, C.E. (1995). Expression and regulation by interferon of a double-stranded-RNA-specific adenosine deaminase from human cells: evidence for two forms of the deaminase. Mol Cell Biol 15, 5376–5388. 10.1128/MCB.15.10.5376.

Peng, Z., Cheng, Y., Tan, B.C., Kang, L., Tian, Z., Zhu, Y., Zhang, W., Liang, Y., Hu, X., Tan, X., et al. (2012). Comprehensive analysis of RNA-Seq data reveals extensive RNA editing in a human transcriptome. Nat Biotechnol 30, 253–260. 10.1038/nbt.2122.

Pestal, K., Funk, C.C., Snyder, J.M., Price, N.D., Treuting, P.M., and Stetson, D.B. (2015). Isoforms of RNA-Editing Enzyme ADAR1 Independently Control Nucleic Acid Sensor MDA5- Driven Autoimmunity and Multi-organ Development. Immunity 43, 933–944. 10.1016/j.immuni.2015.11.001.

Picardi, E., D’Erchia, A.M., Lo Giudice, C., and Pesole, G. (2017). REDIportal: a comprehensive database of A-to-I RNA editing events in humans. Nucleic Acids Res 45, D750–D757. 10.1093/nar/gkw767.

Picardi, E., and Pesole, G. (2013). REDItools: high-throughput RNA editing detection made easy. Bioinformatics 29, 1813–1814. 10.1093/bioinformatics/btt287.

Placido, D., Brown, B.A., 2nd, Lowenhaupt, K., Rich, A., and Athanasiadis, A. (2007). A left-handed RNA double helix bound by the Z alpha domain of the RNA-editing enzyme ADAR1. Structure 15, 395–404. 10.1016/j.str.2007.03.001.

Poulsen, H., Nilsson, J., Damgaard, C.K., Egebjerg, J., and Kjems, J. (2001). CRM1 mediates the export of ADAR1 through a nuclear export signal within the Z-DNA binding domain. Mol Cell Biol 21, 7862–7871. 10.1128/MCB.21.22.7862-7871.2001.

Quinlan, A.R., and Hall, I.M. (2010). BEDTools: a flexible suite of utilities for comparing genomic features. Bioinformatics 26, 841–842. 10.1093/bioinformatics/btq033.

Ricci, E.P., Kucukural, A., Cenik, C., Mercier, B.C., Singh, G., Heyer, E.E., Ashar-Patel, A., Peng, L., and Moore, M.J. (2014). Staufen1 senses overall transcript secondary structure to regulate translation. Nat Struct Mol Biol 21, 26–35. 10.1038/nsmb.2739.

Rice, G.I., Kasher, P.R., Forte, G.M., Mannion, N.M., Greenwood, S.M., Szynkiewicz, M., Dickerson, J.E., Bhaskar, S.S., Zampini, M., Briggs, T.A., et al. (2012). Mutations in ADAR1 cause Aicardi-Goutieres syndrome associated with a type I interferon signature. Nat Genet 44, 1243–1248. 10.1038/ng.2414.

Rice, G.I., Kitabayashi, N., Barth, M., Briggs, T.A., Burton, A.C.E., Carpanelli, M.L., Cerisola, A.M., Colson, C., Dale, R.C., Danti, F.R., et al. (2017). Genetic, Phenotypic, and Interferon Biomarker Status in ADAR1-Related Neurological Disease. Neuropediatrics 48, 166–184. 10.1055/s-0037-1601449.

Ryter, J.M., and Schultz, S.C. (1998). Molecular basis of double-stranded RNA-protein interactions: structure of a dsRNA-binding domain complexed with dsRNA. Embo J 17, 7505–7513.

Schade, M., Turner, C.J., Lowenhaupt, K., Rich, A., and Herbert, A. (1999). Structure-function analysis of the Z-DNA-binding domain Zalpha of dsRNA adenosine deaminase type I reveals similarity to the (alpha + beta) family of helix-turn-helix proteins. Embo J 18, 470–479.

Schwartz, T., Rould, M.A., Lowenhaupt, K., Herbert, A., and Rich, A. (1999). Crystal structure of the Zalpha domain of the human editing enzyme ADAR1 bound to left-handed Z-DNA. Science 284, 1841–1845. 10.1126/science.284.5421.1841.

Shigeyasu, K., Okugawa, Y., Toden, S., Miyoshi, J., Toiyama, Y., Nagasaka, T., Takahashi, N., Kusunoki, M., Takayama, T., Yamada, Y., et al. (2018). AZIN1 RNA editing confers cancer stemness and enhances oncogenic potential in colorectal cancer. JCI Insight 3. 10.1172/jci.insight.99976.

Soker, S., Takashima, S., Miao, H.Q., Neufeld, G., and Klagsbrun, M. (1998). Neuropilin-1 is expressed by endothelial and tumor cells as an isoform-specific receptor for vascular endothelial growth factor. Cell 92, 735–745. 10.1016/s0092-8674(00)81402-6.

Strehblow, A., Hallegger, M., and Jantsch, M.F. (2002). Nucleocytoplasmic distribution of human RNA-editing enzyme ADAR1 is modulated by double-stranded RNA-binding domains, a leucine-rich export signal, and a putative dimerization domain. Mol Biol Cell 13, 3822–3835. 10.1091/mbc.E02-03-0161.

Sun, T., Yu, Y., Wu, X., Acevedo, A., Luo, J.D., Wang, J., Schneider, W.M., Hurwitz, B., Rosenberg, B.R., Chung, H., and Rice, C.M. (2021). Decoupling expression and editing preferences of ADAR1 p150 and p110 isoforms. Proc Natl Acad Sci U S A 118. 10.1073/pnas.2021757118.

Tang, Q., Rigby, R.E., Young, G.R., Hvidt, A.K., Davis, T., Tan, T.K., Bridgeman, A., Townsend, A.R., Kassiotis, G., and Rehwinkel, J. (2021). Adenosine-to-inosine editing of endogenous Z-form RNA by the deaminase ADAR1 prevents spontaneous MAVS-dependent type I interferon responses. Immunity 54, 1961–1975 e1965. 10.1016/j.immuni.2021.08.011.

Wang, F., He, J., Liu, S., Gao, A., Yang, L., Sun, G., Ding, W., Li, C.Y., Gou, F., He, M., et al. (2021). Comprehensive RNA editome reveals that edited Azin1 partners with DDX1 to enable hematopoietic stem cell differentiation. Blood. 10.1182/blood.2021011314.

Wang, Q., Miyakoda, M., Yang, W., Khillan, J., Stachura, D.L., Weiss, M.J., and Nishikura, K. (2004). Stress-induced apoptosis associated with null mutation of ADAR1 RNA editing deaminase gene. J Biol Chem 279, 4952–4961. 10.1074/jbc.M310162200.

Wheeler, E.C., Van Nostrand, E.L., and Yeo, G.W. (2018). Advances and challenges in the detection of transcriptome-wide protein-RNA interactions. Wiley Interdiscip Rev RNA 9. 10.1002/wrna.1436.

Zambelli, F., and Pavesi, G. (2015). RIP-Seq data analysis to determine RNA-protein associations. Methods Mol Biol 1269, 293–303. 10.1007/978-1-4939-2291-8_18.

Zhao, H., Sun, Z., Wang, J., Huang, H., Kocher, J.P., and Wang, L. (2014). CrossMap: a versatile tool for coordinate conversion between genome assemblies. Bioinformatics 30, 1006–1007. 10.1093/bioinformatics/btt730.

Zhao, Z., Zuber, J., Diaz-Flores, E., Lintault, L., Kogan, S.C., Shannon, K., and Lowe, S.W. (2010). p53 loss promotes acute myeloid leukemia by enabling aberrant self-renewal. Genes Dev 24, 1389–1402. 10.1101/gad.1940710.

